# A coordinated and reproducible multicellular response to AD pathology

**DOI:** 10.64898/2026.10.19.759610

**Authors:** Gabriele Vilkaite, Emma Volcov, Lijun An, Yu Xiao, Ines Hristovska, Camila Rosat Consiglio, Alexa Pichet-Binette, Niklas Mattsson-Carlgren, Chris Gaiteri, Jacob W. Vogel

## Abstract

While single-cell measurements of the Alzheimer’s disease (AD) brain allow investigation of vulnerable cells in unprecedented depth, reported findings are heterogeneous, mostly descriptive and often ignore interdependence of cellular responses. Operationalization of these findings requires a multi-cellular model of AD that is robust and reproducible. Here, we utilize a coordinated multicellular community approach across 2.3 million nuclei from 427 ROSMAP donors to identify robust multicellular axes associated with AD pathology and sex. The AD signature comprised profound alteration to RORB-GABRG excitatory neurons and disrupted oligodendrocyte precursor cell signaling. A separate set of AD-related cell-type specific molecular processes with strong sexual dimorphism also emerged. Importantly, these signatures developed and ROSMAP predicted neuropathology and sex, respectively, in a completely independent dataset (SEA-AD). These findings altogether summarize a highly consistent multi-cellular response to AD pathology in the human brain, creating a baseline cellular model to which perturbation and treatment experiments can be compared.

## INTRODUCTION

Alzheimer’s disease (AD) is the most common cause of dementia, characterized by the progressive accumulation of β-amyloid plaques, and neurofibrillary tau tangles. However these hallmark pathologies do not by themselves explain the broader cellular and molecular changes accompanying the disease progression or the variability in cognitive impairment. Recent human transcriptomics atlas studies increasingly support a view where amyloid and tau unfold alongside broader disruptions in synaptic function, neuroinflammation and glial dysfunctions, suggesting that AD reflects a failure of a brain ecosystem as a whole rather than a purely cell-autonomous pathology accumulation ^1–4^. Single-cell and single-nucleus RNA-sequencing studies have further shown that these changes are highly cell type-and cell subtype-specific, affecting excitatory and inhibitory neurons, astrocytes, microglia, oligodendrocytes, OPCs, and vascular-associated cells in distinct ways ^4–6^. Selective vulnerability and resilience also vary across cellular populations, for example, RORB-expressing excitatory neurons and specific inhibitory neuron subtypes showing distinct patterns of disease-associated alteration across AD stages ^7–9^.

At the same time, increasing evidence suggests that the distinct cellular and molecular changes identified across studies may represent components of a broader coordinated multicellular response to AD pathology. In aged human cortex, neuronal, glial and endothelial populations have been shown to co-vary as coordinated multicellular communities associated with AD pathology and cognitive decline ^10^. Trajectory-based analyses have similarly identified coordinated cellular changes across disease progression, particularly among glial support cells ^8,11^. These coordinated responses may also be shaped by communication between neighboring neuronal and glial cells within the same tissue environment ^10,12–14^. To provide clear and robust summarization of single cell AD data, we therefore utilise a coordinated multicellular community (CMC) approach to single cell Alzheimer’s data, emphasizing coherent transcriptional changes across multiple cell types.

However, comparing multicellular structures across studies remains technically difficult. Differences in cohorts, brain regions, disease and pathology measures, and analytical frameworks limit direct comparison of the multicellular patterns reported across studies. Although these studies identify broadly related disease processes, they often emphasize different cellular populations and capture different components of the wider multicellular response. Additionally, benchmarking studies have shown that analytical choices across single-cell RNA-sequencing workflows, such as normalization, filtering and clustering, can alter detected expression differences and cell type assignments, substantially influencing downstream findings and biological interpretation ^15,16^. These sources of variation make it difficult to determine whether differences between studies reflect genuine biological heterogeneity or analytical instability. Identifying coordinated changes that remain robust and reproducible across independent cohorts is therefore important for moving beyond individual descriptions towards a more integrated model of cellular response to AD pathology. There has also been limited exploration as to how these multicellular signatures vary in association with demographic and genetic factors. Sex is particularly relevant, as women are more frequently affected by AD, have been reported to show greater tau burden and faster tau accumulation than men ^17,18^, sex-associated immune and glial differences in AD ^19,20^.

To address these gaps, we applied the CMC framework to single-nucleus transcriptomic data from the dorsolateral prefrontal cortex of 427 donors from the Religious Orders Study and Memory and Aging Project (ROSMAP) to identify a robust and reproducible multicellular representation of AD-associated transcriptional variation and determine whether the resulting gene modules formed coordinated axes associated with neuropathology. To ensure robustness, we systematically evaluated 432 preprocessing and clustering parameter combinations to identify a reproducible and biologically interpretable representation of disease-associated transcriptional variation, and tested its reproducibility in the independent SEA-AD cohort. We next examined how these coordinated multicellular responses change across increasing pathology, relate to cognitive decline, and are connected through predicted cell-cell communication. Results from these approaches support a multicellular model of AD, in which disease progression reflects coordinated changes across interacting brain cell populations rather than isolated alterations within individual cell types.

## RESULTS

### Overview of the study

We developed a modular analytical pipeline and applied it to approximately 2.3 million nuclei from the dorsolateral prefrontal cortex of 427 ROSMAP donors, including 146 cognitively unimpaired individuals, 110 with mild cognitive impairment, and 171 with dementia ^4^ to identify coordinated multicellular transcriptional changes robustly associated with AD-related phenotypes (Fig. 1A). We first preprocessed the data at the single-cell level and constructed donor-and cell subtype-specific metacells, groups of transcriptionally similar cells aggregated to reduce data sparsity and technical noise. Metacells were then split into training, validation and test sets, whilst ensuring that metacells from the same donor were assigned to the same set. Within each cell subtype, genes were clustered into co-expression modules, and linear mixed-effects modelling was used to identify modules associated with AD-related pathological and clinical phenotypes, including global AD pathology burden, amyloid and tau burden, neuropathological staging, vascular and co-pathology measures, APOE genotype and clinical diagnosis. Modules that showed robust associations across training and validation datasets were retained as AD-associated gene modules (AGMs). These AGMs were used in partial least squares (PLS) regression to identify multivariate AGM-phenotype relationships, and were further characterized through other downstream analyses. Findings were replicated in the independent SEA-AD snRNA-seq cohort ^8^.

**Fig. 1.**
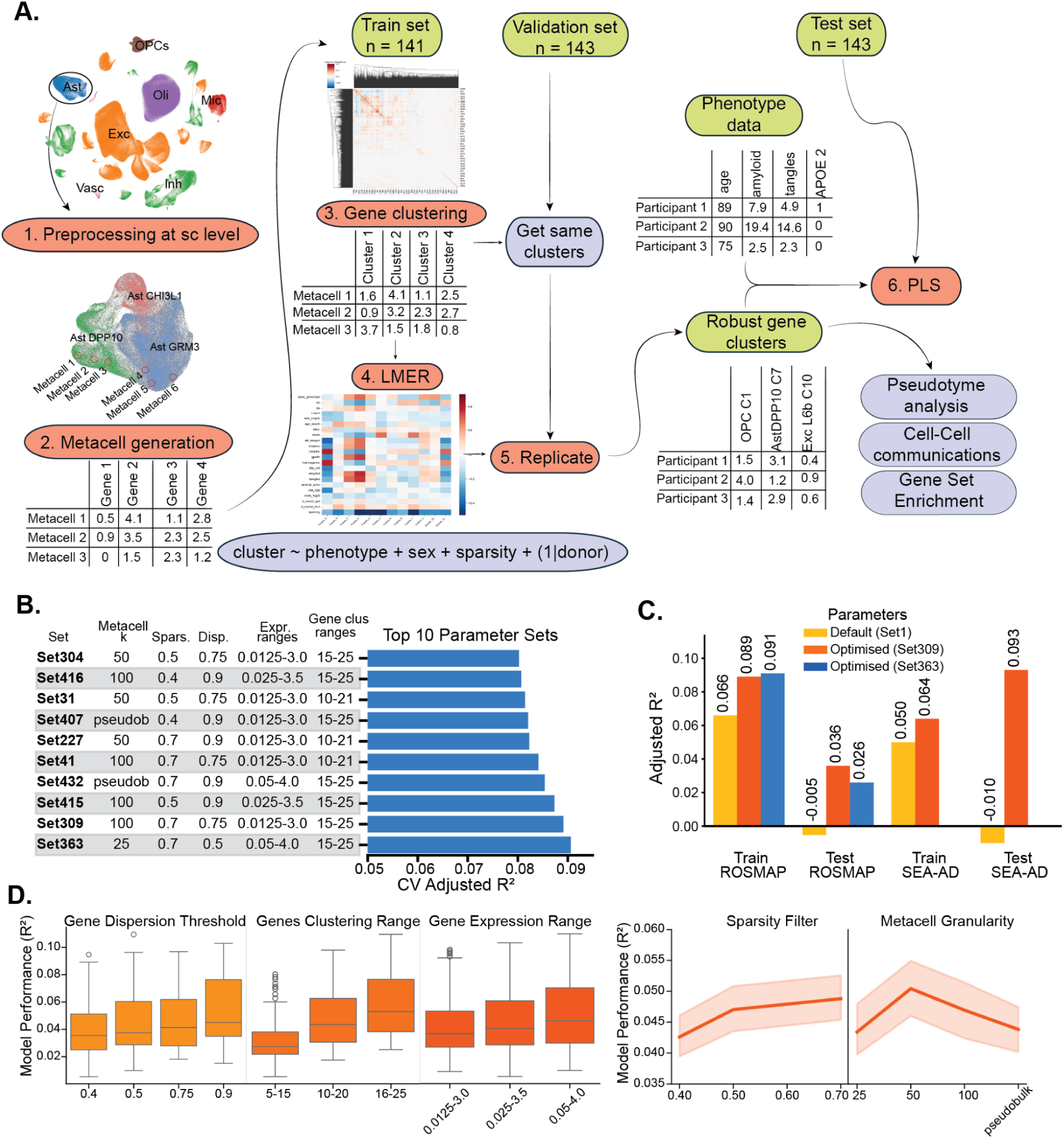
Pipeline workflow optimisation. (A) Pipeline illustrating the analytical workflow. Single-nuclei RNA-seq data from dorsolateral prefrontal cortex (427 donors, ∼2.3 million nuclei) was preprocessed, and cells were clustered into metacells via k-means. Data was split into three parts: train set, validation set and test set. The train set was used for gene clustering into networks and used in the LMER model to select AD-associated gene co-expression clusters. The validation set was used for replication to identify and robust AD-associated gene co-expression modules to be used in Partial Least Squares (PLS) regression analysis and other downstream analyses. **(B) Top 10 parameter sets ranked** by PLS cross-validated model’s adjusted-R2 performance. **(C) Barplot comparison of optimized parameters (Set 363 and Set 309) versus default settings (Set 1),** showing cross-validated adjusted-R2 value in training set and predictive adjusted-R2 value in test sets. **(D) Effects of individual preprocessing parameters** on model performance of all runs, including gene dispersion threshold, gene clustering range, gene expression range, metacell granularity, and sparsity filtering.

### A systematic optimisation of snRNA-seq data preprocessing improves robustness of snRNA-phenotypes associations

Preprocessing and clustering parameter choices can strongly influence the detection of AD-associated gene modules, including how cells are aggregated, which genes are retained, and how co-expression modules are defined. Since our goal was to develop a robust cellular signature of AD, we systematically optimised key preprocessing and clustering parameters using a grid search of 432 predefined parameter combinations. The parameter grid included metacell granularity, gene expression thresholds, gene dispersion thresholds, sparsity filters, and gene clustering ranges (Table 1), with the model performance evaluated using adjusted cross-validated R2 (Fig. 1A).

**Table 1.** Parameter values evaluated during pipeline optimization. Parameter combinations were evaluated using R2 values of the downstream PLS model. The set with the highest training set R2 value was chosen.

|  |  |  |  |  |
| --- | --- | --- | --- | --- |
| Gene Expression Dispersion Threshold | 0.4 | 0.5 | 0.75 | 0.9 |
| Metacell $k$ | 25 | 50 | 100 | pseudobulk |
| Min and Max Gene expression cutoffs | 0.0125 - 3.0 | 0.025 - 3.5 | 0.05 - 4.0 |  |
| Metacell sparsity thresholds | 0.4 | 0.5 | 0.7 |  |
| Gene clustering $k$ ranges | 5-15 | 10-20 | 15-25 | |

Next, we ranked all pipeline runs by the adjusted cross-validated R2 of the resulting PLS models, evaluating how well they summarised the main relationships between AD-associated gene module expression and phenotypes. The two best performing parameter sets, Set363 and Set309, achieved similar model performance, (Fig. 1B; Supplementary Table 1). However, Set309 provided a more parsimonious and biologically interpretable model using two PLS latent components. In contrast, Set363 required five components to explain a similar amount of variance, making the model more difficult to interpret.

To assess whether optimisation improved performance beyond standard preprocessing choices, we compared the two best-performing parameter sets, Set 309 and Set 363, with the default parameter set, Set 1, which represented Scanpy package’s default preprocessing parameters and the reference clustering configuration defined prior to optimisation (Fig. 1C). In the training dataset, optimisation increased adjusted cross-validated R2 from 0.066 in Set 1 to 0.089 in Set 309 and 0.091 in Set 363, corresponding to an approximately 33% improvement. This improvement generalised to the independent ROSMAP test set, where adjusted R2 increased from-0.005 in Set 1 to 0.036 in Set 309 and 0.026 in Set 363. To further evaluate external generalisability, we applied the same analytical framework to the SEA-AD prefrontal cortex cohort by reconstructing ROSMAP-derived AD-associated gene modules in SEA-AD. Consistent with the ROSMAP results, Set 309 showed improved performance in both train and test sets compared with the default parameter set. Overall, Set 309 provided the best balance between predictive performance, generalisability, and interpretability, and was therefore selected for downstream analyses.

Notably, many of the top performing parameter sets had similar R2 values and converged on similar parameter ranges. Aggregating performance across all runs revealed systematic effects of preprocessing choices (Fig. 1D). Higher gene clustering ranges, and thus smaller, finer gene co-expression modules, were associated with improved R2 value of PLS models (Fig. 1D). Higher gene dispersion thresholds, which restrict highly variable genes (HVG) selection to only the most variable genes, also improved model performance. This suggests that focusing on only the strongest and most variable transcriptional signals enhances the stability and predictive power of the downstream models. Similarly, more granular metacells tended to increase performance, whereas extreme settings such as pseudobulk aggregation of each cell subtype per each donor slightly reduced it.

### Identification of robust AD-associated gene modules

To identify gene modules robustly associated with AD-related phenotypes, we next performed linear mixed-effects regression (LMER) analyses across all gene modules and neuropathological and clinical phenotypes (Fig. 2A). Module-phenotype associations were first tested in the training subset, and significant module-phenotype associations (FDR < 0.05) were further re-tested in a validation subset using nominal (p<0.05) significance thresholds (Fig. 2A; Supplementary Table 2). To prioritize reproducible associations and reduce false-positive findings from high-dimensional testing, we retained only modules that were significant in both datasets and showed consistent effect directionality. These reproducible modules were defined as AD-associated gene modules (AGMs).

**Fig. 2.**
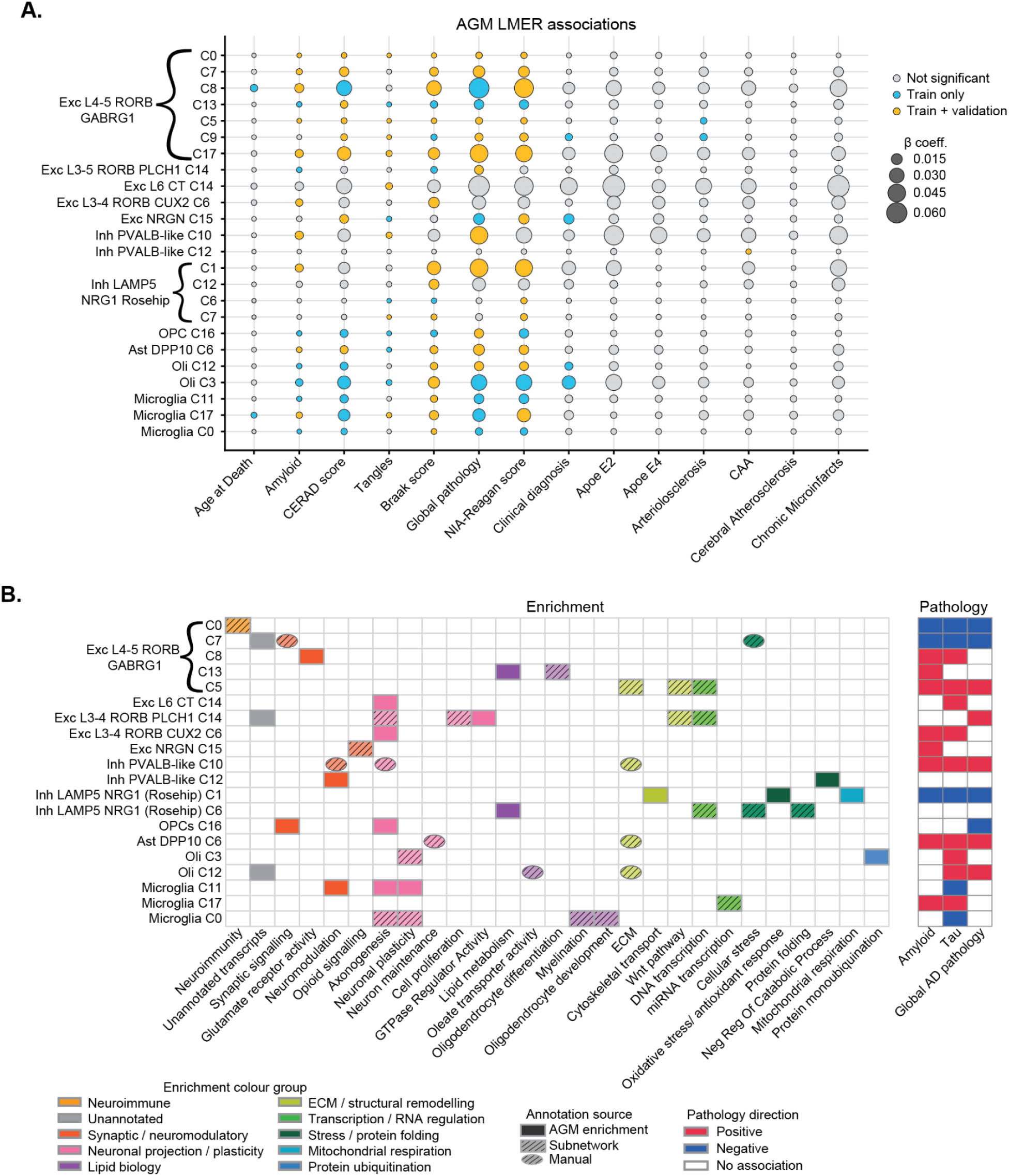
Reproducible associations between AD-associated gene modules and clinical or neuropathological traits. (A) Dot plot of LMER associations between AD-associated gene modules (AGMs) and clinical or neuropathological phenotypes. Dot size represents the absolute LMER β coefficient from the training set. Colours indicate association status: grey, not significant; blue, significant in the training set only after FDR correction; pink, significant in both the training set and validation set, using nominal significance in validation. Only AGMs demonstrating robust AD-associated effects are shown. **(B) Functional enrichment and pathology associations of AD-associated gene modules.** The heatmap (left) shows broad functional enrichment categories associated with each AGM. Colours indicate the functional group of the enrichment term. Annotation style indicates the level of confidence: solid-filled cells represent significant AGM-level enrichment (FDR<0.05), hatched cells represent enrichment of subnetworks within AGMs (FDR<0.05), and circles indicate weaker, manual annotations. The pathology panel (right) shows directions of association between AGMs and amyloid (amyloid and CERAD), tau (tangles and Braak stages), and global AD pathology burden. Only AGMs that had significant functional enrichment are shown.

We identified 24 AGMs out of 533 tested gene modules, corresponding to approximately 4.5% of all modules evaluated, indicating that only a restricted subset of transcriptional programs demonstrated reproducible associations with AD-related phenotypes. The 24 AGMs were distributed across multiple major cell classes, but they were not evenly represented across cell types. Eleven AGMs mapped to excitatory neuronal populations, including seven distinct AGMs from the Exc L4-5 RORB GABRG1 subtype alone. Six AGMs mapped to inhibitory neurons, including four from the Inh LAMP5 NRG1 (Rosehip) population and two from Inh PVALB-like interneurons. The remaining AGMs comprised one astrocytic module, one OPC module, two oligodendrocyte modules, and three microglial modules. Replicated associations were primarily found among neuropathological traits such as Braak score, which had 15 replicable AGMs associated with it, global AD pathology burden (See Methods) (11 AGMs), NIA-Reagan score (13 AGMs), amyloid plaques (10 AGMs) and tau tangles (8 AGMs associated). Notably, AGMs were found almost exclusively for AD-associated phenotypes. The lone exception was Inhibitory PVALB-like gene cluster 12, which was associated only with CAA.

To interpret the biological functions represented by the AGMs, we next annotated the modules using complementary enrichment strategies (Fig. 2B, Supplementary Table 3-4). First, each full AGM gene list was tested for over-representation of GO Biological Processes and KEGG terms, identifying significant functional enrichment for 12 AGMs (FDR < 0.05). To more thoroughly characterize AGMs, we additionally used STRING network analysis to identify connected gene subnetworks within each AGM and tested them for functional enrichment (FDR<0.05). This allowed us to detect biologically coherent gene groups embedded within larger AGMs, even when the full module did not show significant enrichment. Finally, exploratory annotations were added for modules with recurrent genes or recognizable functional themes that did not reach statistical significance in the enrichment analyses. These three annotation levels are shown separately in Fig. 2B.

Together, these approaches provided functional annotations for 20 of the 24 AGMs, whereas four modules showed no clear enrichment and could not be readily interpreted. Annotated AGMs captured a wide range of biological processes, including synaptic signaling, neuronal plasticity and axonogenesis, extracellular matrix (ECM) organisation as well as cellular stress and protein-folding pathways. We also tested the AGMs for enrichment in long non-coding RNAs (lncRNAs) and unannotated transcripts using Fisher’s exact test. While none of the AGMs were significantly enriched for lncRNAs (FDR<0.05), several modules were significantly enriched for unannotated transcripts. The limited functional information available for these transcripts may partly explain why some AGMs were more difficult to annotate.

Beyond the presence of these broad functional categories, their distribution across cell types was informative. Axonogenesis, neuronal projection and plasticity-related annotations were present in AGMs from several excitatory, glial and microglial populations, including Exc L6 CT, Exc L3-4 RORB PLCH1, Exc L3-4 RORB CUX2, OPCs, oligodendrocytes and microglia. In contrast, these annotations were absent from the AGMs belonging to Excitatory L4-5 RORB GABRG1 and Inhibitory

LAMP5 NRG1 (Rosehip) neurons, two neuronal populations previously implicated to be vulnerable in AD. Instead, AGMs from these vulnerable populations were mainly associated with synaptic receptor activity, ECM organisation, transcriptional regulation, cellular stress, protein folding and mitochondrial respiration, suggesting a stronger association with synaptic dysfunction and cellular stress rather than axonal or structural plasticity.

Exc L4-5 RORB GABRG1 neurons showed several diverse AGMs, with C0 and C7 negatively associated with AD pathology, whereas C8, C13 and C5 were positively associated with at least one AD hallmark pathology (Fig 2B, right panel). C0 contained a 53-gene STRING subnetwork linked to neuroimmune signalling (Fig 2B; Supplementary Fig. 1A). C7, while harder to characterise due to enrichment in unannotated transcripts, included several known synaptic and stress-related genes, including *GRIN3A*, *SNCG*, *LAMP5*, *CRYAB*, *CIRBP*, *ROMO1* and *SH3BGRL3*. In contrast, C8 showed a clear glutamatergic profile, with enrichment in terms associated to modulation of synaptic transmission and G protein-coupled glutamate receptor signaling, and contained genes including *GRIK2*, *CNTNAP*, *CNTN* and *CDH12/18* and GRM family members (Fig 2B; Supplementary Fig. 1B). The positive association of this glutamatergic receptor and synaptic signalling module with pathology suggests that increase in AD pathology is also associated with a shift in excitatory synaptic signaling within Exc L4-5 RORB GABRG1 neurons.

Some AGMs also pointed to more pathology-specific cellular states. Inhibitory PVALB-like C12 was selectively associated with cerebral amyloid angiopathy, rather than broader AD pathology measures. It showed significant enrichment for neuromodulatory/neuropeptide activity, including genes such as *NPY*, *PNOC* and *VIP*, while also containing vascular-associated genes such as *TIMP1*, *MET*, *RASIP1* and *MFSD2A*. Together, its CAA-specific association and neuromodulatory/vascular profile suggest that C12 may capture a PVALB-like interneuron state linking neurovascular coupling with vascular amyloid pathology. OPC C16 was negatively associated with global AD pathology and enriched for axonogenesis and synaptic signalling terms, including chemical synaptic transmission and voltage-gated cation channel activity, with genes such as *GABRA4*, *GABRG2* and *GABRG3* (Fig 2B; Supplementary Fig. 1D). This suggests that increasing pathology is accompanied by loss of OPC functions related to neuronal and synapse sensing and circuit maintenance.

Additionally, the three microglial AGMs also showed distinct functional profiles. Microglia C0 and C11 were negatively associated with pathology and contained annotations related to neuronal maintenance, including axonogenesis, neuromodulation, neuronal plasticity, and, within specific subnetworks, myelination-related processes. Their negative associations with pathology suggest that these supportive transcriptional programs become less prominent with increasing pathological burden, consistent with the negatively associated neuroimmune-signalling module in Exc L4-5 RORB GABRG1 neurons C0. Overall, the functional diversity and contrasting pathology associations of the AGMs indicate that AD is accompanied not by a single uniform transcriptional response, but by multiple distinct neuronal and glial biological processes that are differentially altered across disease pathology.

### PLS identifies distinct pathology-and sex-associated multicellular transcriptional axes

After identifying disease-associated transcriptional changes, we next sought to understand whether or not these changes were independently associated with AD pathology. To do this, we applied Partial Least Squares (PLS) regression to model the multivariate relationship between donor-level gene module expression of these 24 AGMs and AD-associated neuropathological and clinical phenotypes. Adjusted cross-validated model performance was highest for a two-component model (total multi-dataset covariance R2=0.09; Fig. 3A). Permutation testing confirmed that both components captured non-random structure (both PLS Component 1 and Component 2 empirical p=0.001; Fig. 3B), supporting the selection of a two-component solution.

**Fig. 3.**
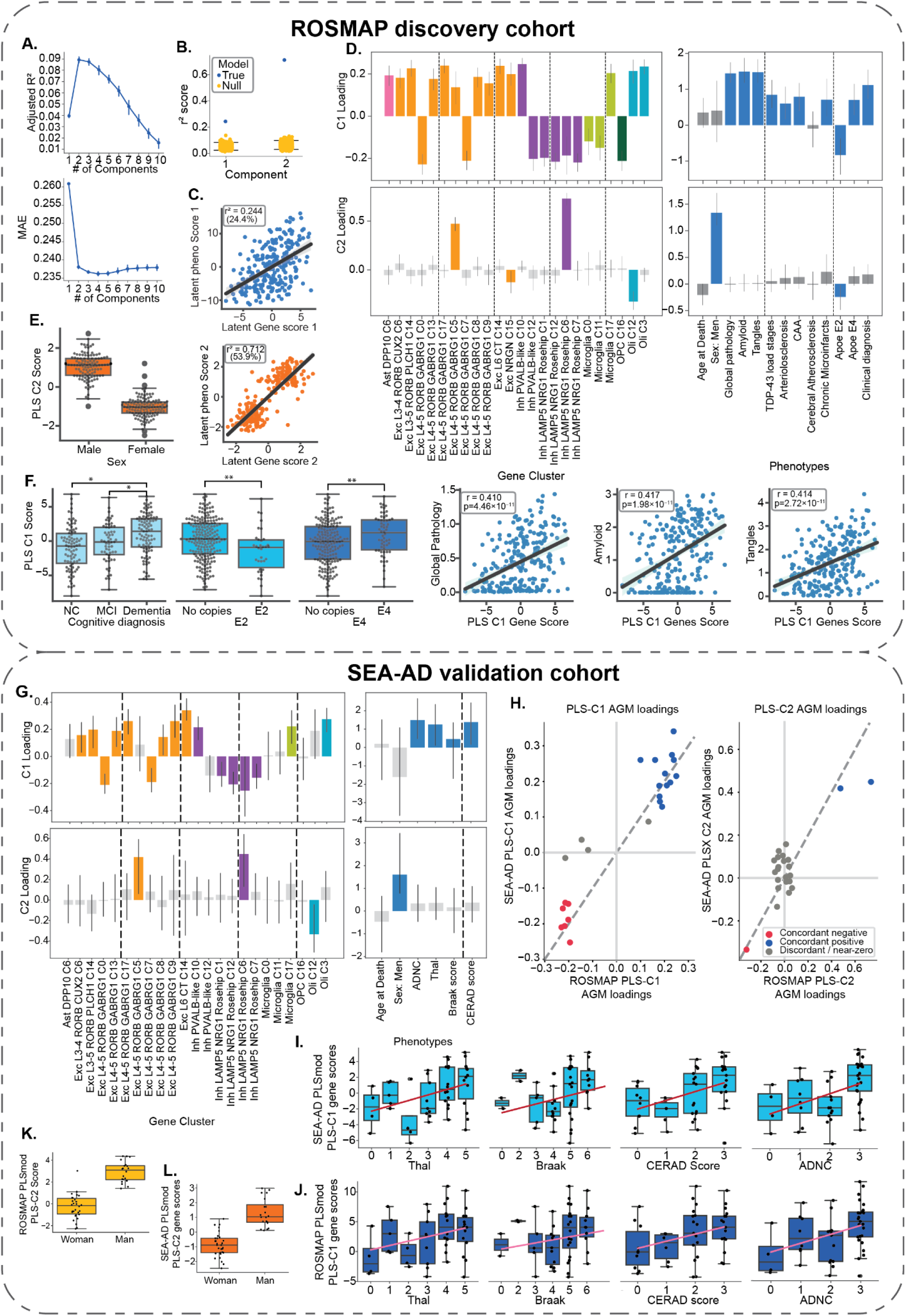
Partial least squares (PLS) modelling of coordinated AGM variation in the ROSMAP discovery cohort and replication in the SEA-AD validation cohort. (A) PLS model performance in ROSMAP. Adjusted cross-validated R2 (top) and mean absolute error (MAE; bottom) across models with increasing numbers of PLS components. (B) Permutation test comparing the observed model R2 (blue) with the null distributions (yellow) for the component. (C) Scatterplot of latent gene scores versus latent phenotype scores for components 1 (PLS-C1) and 2 (PLS-C2), show the shared covariance captured. (D) ROSMAP PLS component loadings. AGM loadings (left) and phenotype loadings (right) for PLS-C1 (top) and PLS-C2 (bottom). Significant AGM loadings (p < 0.05) are colour-coded by major cell type, while non-significant loadings are shown in grey. (E) PLS C2 gene scores stratified by sex in the ROSMAP cohort. (F) Associations between ROSMAP PLS-C1 gene scores and clinical, genetic, and neuropathological phenotypes, including cognitive diagnosis, APOE E2 and E4 carrier status, global AD pathology, amyloid, and tangle burden. (G) De novo PLS model fitted to the SEA-AD validation cohort, showing AGM (left) and phenotype (right) loadings for PLS-C1 (top) and PLS-C2 (bottom). (H) Concordance of AGM loadings between the ROSMAP and de novo SEA-AD PLS models for C1 (left) and C2 (right). Each point represents a matched AGM; blue and red indicate concordant positive and negative loadings, respectively, while grey indicates discordant or near-zero loadings. (I) SEA-AD de novo PLS-C1 gene scores across neuropathological measures: Thal phase, Braak stage, CERAD score, and AD neuropathological change (ADNC). (J) ROSMAP PLS-C1 gene scores after projection of the ROSMAP model into SEA-AD, shown across the same neuropathological measures. (K) SEA-AD de novo PLS-C2 gene scores stratified by sex. (L) ROSMAP PLS-C2 gene scores after projection of the ROSMAP model into SEA-AD stratified by sex.

PLS Component 1 (PLS-C1) was associated with a broad pathology-linked latent axis, accounting for 24.7% of the shared covariance between the AGM expression and the phenotypes, and was driven by contributions from all selected AGMs (Fig. 3C, Supplementary Table 5). On the phenotype side, this component loaded positively with higher global AD pathology, more amyloid plaques and neurofibrillary tau tangles, higher TDP-43 load stages, more severe arteriolosclerosis, cerebral amyloid angiopathy, and chronic microinfarcts, and more advanced clinical diagnosis. APOE E2 loaded negatively, with APOE E2 carriage associated with lower component scores (Fig. 3D).

To further characterize the pathology-associated PLS-C1 axis, we evaluated associations between this multicellular transcriptional signature and AD-associated phenotypes. Consistent with the phenotype loadings, higher PLS-C1 gene scores were associated with greater neuropathological burden, including higher global AD pathology burden, amyloid, and tau tangles (Fig. 3F; Supplementary Fig. 2). Similar positive associations were also observed across regional measures of amyloid, neuritic and dense plaques, and neurofibrillary tangles, indicating that the PLS-C1 association was not driven by pathology in a single brain region, but reflected a more broad AD pathology signature (Supplementary Fig. 3A-D). PLS-C1 gene scores were also higher in donors with dementia than in cognitively normal donors or those with mild cognitive impairment, and showed opposing associations with APOE genotype, lower scores in APOE E2 carriers and higher scores in APOE E4 carriers. Together, these findings support PLS-C1 as a coordinated multicellular transcriptional axis associated with increasing AD pathology and more advanced clinical disease.

Component 2 (PLS-C2), on the other hand, captured a distinct sex-associated axis. Although it explained 53.8% of leftover covariance after PLS-C1, only four AGMs loaded significantly onto this component (Fig. 3C, Supplementary Table 5). The dominant phenotype loading on Component 2 was sex, showing positive associations with men and a weaker negative loading for APOE E2. Component 2 gene scores were also strongly separated by sex, with men showing much higher gene scores than women (Fig. 3E). Thus, the two-component PLS solution separated a broadly distributed pathology-associated multicellular program from a more focused sex-linked transcriptional signal.

### The ROSMAP pathology and sex-associated PLS components replicate in SEA-AD cohort

We next tested whether the pathology and sex-associated PLS components identified in the ROSMAP cohort could be reproduced in SEA-AD, an independent snRNA-seq prefrontal cortex dataset with similar AD-related phenotypes. AGMs, previously defined in ROSMAP, were reconstructed using the corresponding genes available in SEA-AD (Supplementary Fig. 4C-D). We first fit a de novo PLS model using the reconstructed SEA-AD AGMs. Cross-validation supported a two-component solution, and permutation testing indicated that the selected components captured non-random structure (Supplementary Fig. 4A). As in ROSMAP, SEA-AD PLS Component 1 was primarily associated with pathology-related phenotypes, whereas Component 2 was aligned with sex (Fig. 3G; Supplementary Fig. 4B). Consistent with the phenotype loadings, higher PLS-C1 gene scores were associated with greater pathology burden measures (Fig. 3I) and PLS-C2 gene scores were higher in men than in women (Fig. 3K). We next compared the loadings of matched AGMs between the ROSMAP and SEA-AD PLS models. AGM loadings showed strong concordance between cohorts for both components (PLS-C1 r=0.94; PLS-C2 r=0.84), with most significantly loading AGMs matching loading directions and indicating that modules contributing most strongly to the component were largely preserved across cohorts (Fig. 3H). Moreover, projecting the ROSMAP-trained PLS model directly onto SEA-AD AGM matrix without retraining reproduced the expected relationships between latent scores, pathology, and sex (Fig. 3J, L). Together, these results indicate that the AGM loading patterns underlying both pathology and sex-associated components are highly consistent between ROSMAP and SEA-AD. This cross-cohort replication indicates that PLS-C1 captures a highly reproducible, shared, dataset-independent transcriptomic trend that robustly tracks AD pathological progression, and PLS-C2 captured consistent, similarly reproducible and pronounced sex-associated differences across both datasets.

### Coordinated multicellular changes track pathology burden and show pathology-dependent associations with cognitive decline

Having established a reproducible, multicellular signature associated with AD pathology, we next sought to characterize the dynamics of this cell network. First, we asked whether the AD-associated modules change in parallel or whether they follow different temporal patterns across the disease continuum. To test this, we modeled AGMs expression as a function of global AD pathology using generalized additive models (GAMs), and grouped the fitted trajectories by spectral clustering. This identified two linked pseudoprogression trajectories (Fig. 5A). One group, comprising 14 AGMs, showed progressive increase in gene expression with increasing global AD pathology, whereas the second group, comprising 10 AGMs, showed a constant decrease in gene expression. Interestingly, modules within each group followed nearly identical trajectories (Supplementary Fig. 5), either rising or falling at similar rates, indicating that their expression changes are likely coordinated in time.

**Figure 5.**
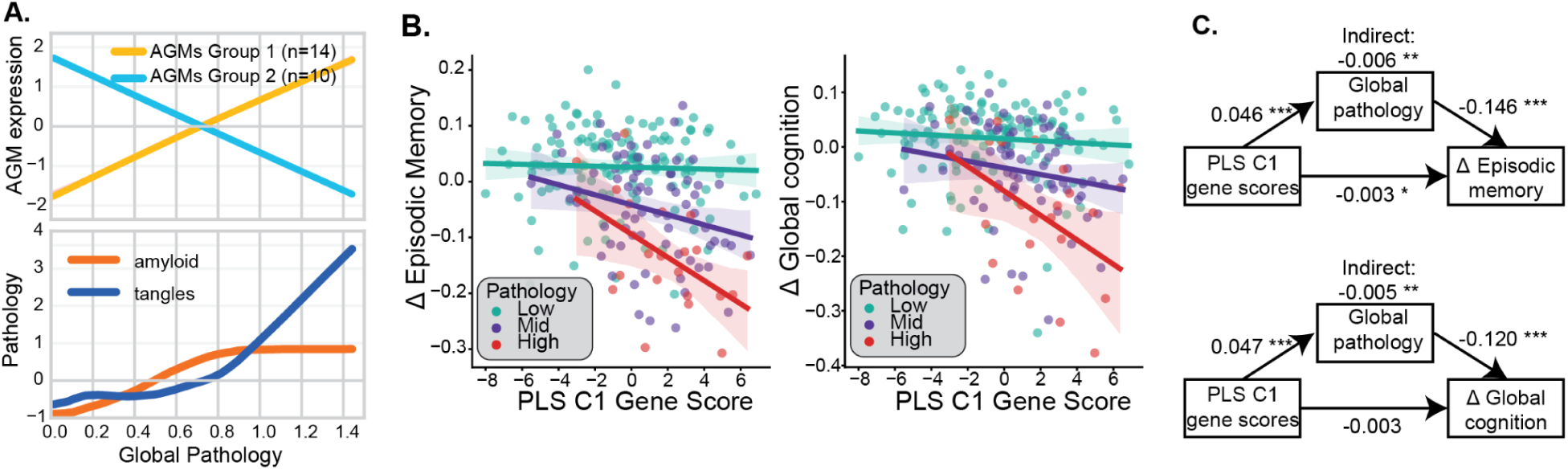
Pseudoprogression, moderation and mediation analyses of AD-associated multicellular transcriptional programs. **(A)** Pseudoprogression trajectories across the log-transformed global AD pathology axis. Top panel - two AGM trajectory groups identified by spectral clustering. Bottom - z-scored amyloid and tangle progression across the global AD pathology axis. **(B)** Moderation analyses showing significant PLS C1-by-global AD pathology interactions for cognitive performance after FDR correction. Regression lines are stratified by pathology groups: low (0–0.5), mid (0.5–1), and high (1–1.5) log-transformed global AD pathology. **(C)** Mediation models for episodic memory, and global cognition. Numbers indicate standardized regression coefficients. Asterisks denote significance (p < 0.05, p < 0.01, p < 0.001).

The increasing trajectory was predominantly composed of excitatory neuronal AGMs, including five of the seven modules from Exc L4-5 RORB GABRG1 neurons, and other excitatory neuron AGMs, together with oligodendrocyte, Ast DPP10, and Inh PVALB-like C10 modules. In contrast, the decreasing trajectory contained several Inh LAMP5 NRG1 (Rosehip) AGMs, the OPC AGM, Exc L4-5 RORB GABRG1 C0 and C7, and the Microglia C0 and C11 modules. These findings reveal a multicellular transcriptional program associated with increasing AD pathology (Supplementary Fig. 5).

Given that PLS-C1 was strongly associated with pathology burden, we next tested whether its relationship with longitudinal cognitive decline differed across levels of global AD pathology. Moderation analysis showed that the association between higher PLS-C1 gene scores and faster decline became stronger as global AD pathology increased for both episodic memory (β=−0.015, 95%CI [−0.025, −0.006], t=−3.16, FDR-p=0.011) and global cognition (β=−0.013, 95%CI [−0.022, −0.003], t=−2.58; FDR-adj p = 0.021; Fig. 5B). At low levels of global AD pathology, PLS-C1 scores showed little association with cognitive change. In contrast, among individuals with high pathology burden, higher PLS-C1 scores were associated with faster antemortem decline in episodic memory and global cognition.

The pathology-dependent association between PLS-C1 and cognitive decline raised the possibility that these multicellular transcriptional changes might mediate the established relationship between AD pathology and cognition. We therefore first tested whether PLS-C1 gene scores mediated the association between global AD pathology and cognitive decline. This model showed no evidence of an indirect effect through PLS-C1 (Supplementary Fig. 6A). We next tested the complementary model, in which global AD pathology mediated the association between PLS-C1 gene scores and cognition. Higher PLS-C1 scores were associated with greater global AD pathology, which in turn was associated with faster decline in episodic memory and global cognition (Fig. 5C, Supplementary Fig. 6B). Significant indirect effects were observed for episodic memory (indirect effect = −0.006, 95%CI[-0.009,-0.004], p<0.001) and global cognition (indirect effect = −0.005, 95%CI[-0.008,-0.003], p<0.001). Global AD pathology accounted for an estimated 66% of the total association between PLS-C1 and episodic memory decline and 58% of the association between PLS-C1 and global cognitive decline (Supplementary Fig. 6C). Together, these results suggest that the multicellular transcriptional process captured by PLS-C1 are at least partially independent of AD neuropathology, but affect cognition mainly through its interaction with neuropathology.

### AD-associated gene modules form a multicellular communication network

The PLS model identified a shared transcriptional pattern across multiple cell populations, and so we next aimed to examine what intercellular crosstalk between different cell types might coordinate the changes identified by the PLS model. To address this, we constructed a predictive cell-cell communications network inferring ligand-receptor, adhesion-adhesion, and gap-junction interactions between AGMs using CellPhoneDb v5.0.0.

The results revealed a dense network of communication including all major cell classes, with predicted ligand-receptor interactions arising from AGMs that increase or decrease with AD pathology, capturing communication changes across disease progression, rather than treating the network as a static disease-state architecture (Fig. 6A, top panel). The observed network contained 125 predicted molecular interactions, including 111 ligand-receptor (LR) interactions distributed across 94 directed AGM-AGM edges, 13 adhesion-adhesion interactions distributed across 12 AGM-AGM edges, and one gap-junction interaction. Because the analysis was restricted to AD-associated AGMs, both genes in each predicted LR pair originated from modules whose expression change was associated with pathology. The amount of cell-cell communication (expressed here as the number of LR pairs) significantly exceeded what would be expected from a similar sized network of random cell subtypes, whereas adhesion-adhesion connectivity remained within the null range (Fig. 6B, Supplementary Table 6).

**Fig. 6.**
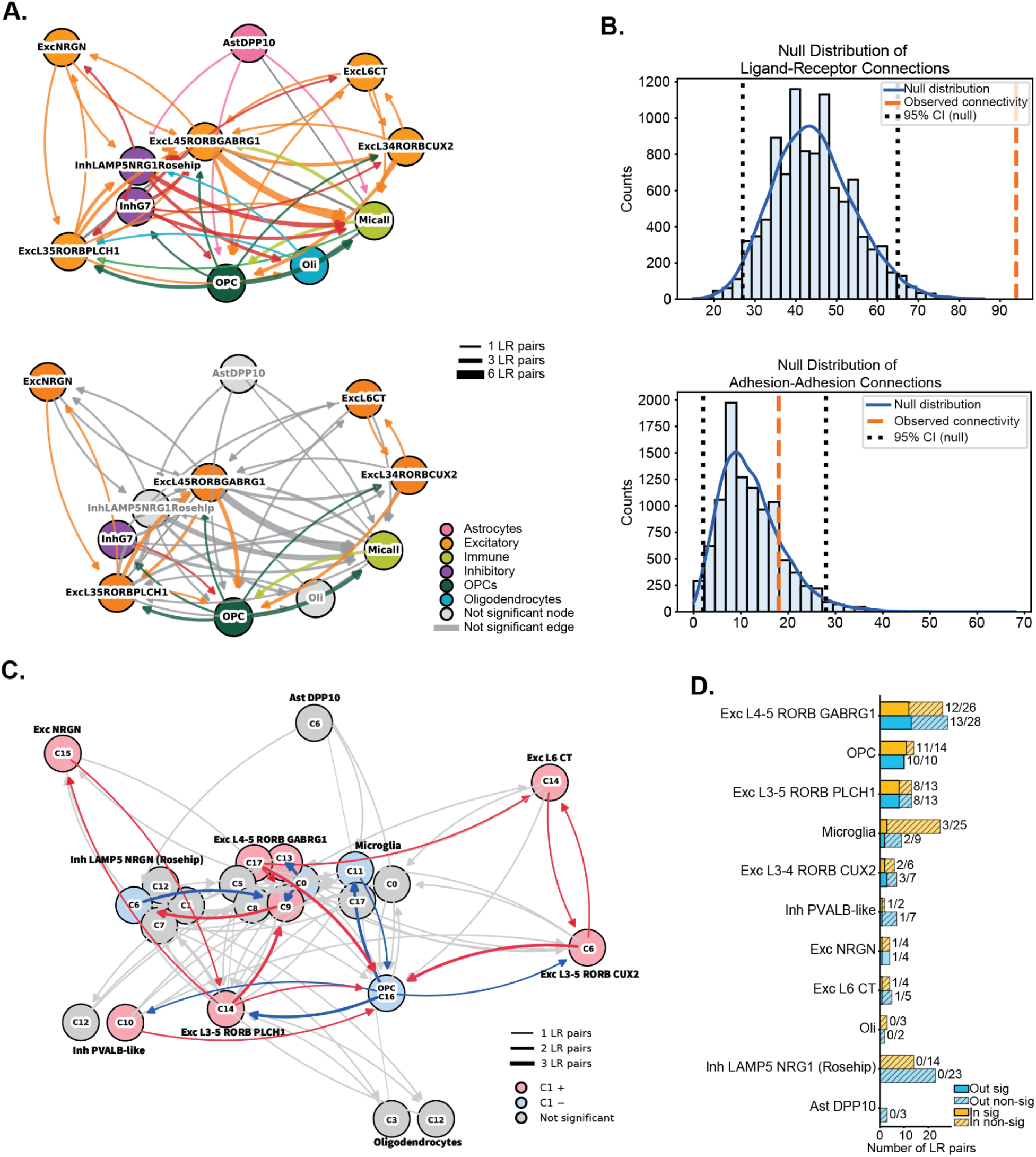
Cell-cell communications network between AGMs. (A) Putative cell-cell communications network between cell subtypes. Nodes represent each cell subtype; colour indicates parent cell type. Arrows represent unidirectional ligand-receptor (LR) connections, with colour corresponding to the node of origin. Grey coloured edges show bidirectional connections, such as gap junctions and adhesion proteins. Edge thickness indicates the number of connections between the nodes. Grey edges show connections which did not pass the permutation test. **Putative cell-cell communications network between cell subtypes.** The same graph as, but this time, nodes and edges that did not survive a null modelling procedure were coloured grey. **(B) CCC null modelling binary edges frequency**, showing number of total edges formed between AGMs, for Ligand-Receptor connections (top) and Adhesion-Adhesion connections (bottom). The red dashed line indicates the observed model’s number or edges, while the blue histogram indicates the distribution of connections between AGMs across null permutations. The black dashed line shows the 95% confidence interval of the null distribution **(C) Putative AGM-AGM network communications.** Nodes and arrows coloured by the PLS Component 1 loading direction (red - positive PLS-C1 loading; blue - negative PLS-C1 loading) AGM nodes that were not significant after null modeling are coloured in pale colours. **(D) Total outgoing and incoming ligand-receptor connections between cell subtypes.** The hashed bar shows the number of significant connections after null modelling, the whole bar shows the total number of LR connections.

We used the null modeling procedure as a conservative measure to prune cell-cell connections that do not exceed chance levels (in terms of number of connections expected between two cell subtypes of similar size and composition; Fig. 6A bottom). The pruned network highlighted several features that were not immediately apparent from the full network. Most notably, OPCs retained reciprocal connections with two excitatory neuronal subtypes, Exc L3-4 RORB CUX2 and Exc L3-5 RORB GABRG1, as well as PVALB-like inhibitory interneurons, and microglia. This indicated that OPC communication remained prominent even after controlling for random network structure. This was also reflected in the sender/receiver summary, where OPCs retained nearly all detected ligand-receptor connections after null modelling, with 10/10 incoming and 11/14 outgoing LR connections passing the null threshold (Fig. 6D). In contrast, several highly connected subtypes in the full network, such as LAMP5 NRG1 Rosehip interneurons, did not retain these connections after null modeling, indicating that high overall connectivity did not necessarily correspond to high-confidence communication.

At the AGM level, OPCs were represented by a single AGM which was negatively associated with AD pathology (Fig. 6C). In the retained network, OPC C16 had reciprocal connections to several neuronal AGMs, many of which were positively associated with pathology. OPC C16 mainly received signals through the receptors *ROBO2*, *ADGRL2*, and *EPHA4*. These interactions were primarily associated with Teneurin, Ephrin, and SLIT/NTRK-like signalling. In the opposite direction, the most frequently detected OPC-expressed ligands were *EFNA5* and *TENM2*, which predominantly interacted with EPHA-family receptors and *ADGRL2* expressed by neuronal, microglial, and OPC AGMs. The predominance of Ephrin-, Teneurin-, and SLIT-related interactions suggests that OPC C16 may contribute to adhesion-and guidance-related communication with neighbouring neuronal and glial populations. This communication profile is consistent with the functional annotation of OPC C16, which also pointed toward neuronal and circuit-associated processes; AGM negative associations thus suggest reduced representation of this OPC-centered communication network as pathology increases.

Worth noting, Exc L4-5 RORB GABRG1 emerged as one of the most connected excitatory subtypes in the retained ligand-receptor network (Fig. 6A, D). However, most significant LR pairs involving this subtype reflected within-subtype communication, out of 13 sent and 12 received LR pairs, 8 originated from within the same subtype, including fibroblast growth factor (FGF), opioid, SLIT and pleiotrophin signaling. The remaining incoming interactions came from Exc L3-5 RORB PLCH1, involving pleiotrophin, opioid, and retinoic-acid-associated signalling, and from OPCs through somatostatin, SLIT and Teneurin signalling (Supplementary Table 6). This suggests that the Exc L4-5 RORB GABRG1 network is dominated by recurrent module-to-module communication, with additional input from neighbouring excitatory and OPC-associated signalling axes. These findings together suggest that AD-associated AGMs are linked by a non-random ligand-receptor network with prominent recurrent excitatory communication and retained OPC-mediated crosstalk.

### AD-associated cellular processes show sexual dimorphism

Sex emerged as the defining phenotype of PLS Component 2, despite these AGMs being selected based on their association with AD-related phenotypes independently of sex. So, we next examined whether individual AGMs exhibited any sex by global AD pathology interactions. Whilst we did not detect any sex * AGM expression interactions on pathology, we did observe several AGMs that were significantly associated with both sex and global AD pathology burden independently (after FDR correction; Fig. 7). Exc L4-5 RORB GABRG1 cluster 5 expression increased with increased pathological burden in both sexes, but had a higher baseline expression in men. Oligodendrocytes cluster 12 likewise showed increased transcription across the global AD pathology burden axis, but was consistently expressed more in women. Lastly, inhibitory neurons LAMP5 NRG1 cluster 6 continuously decreased with greater amounts of pathology, but showed much greater expression levels in men compared to women. Notably, despite allowing for nonlinear trends, these data were best supported by linear models.

**Fig. 7.**
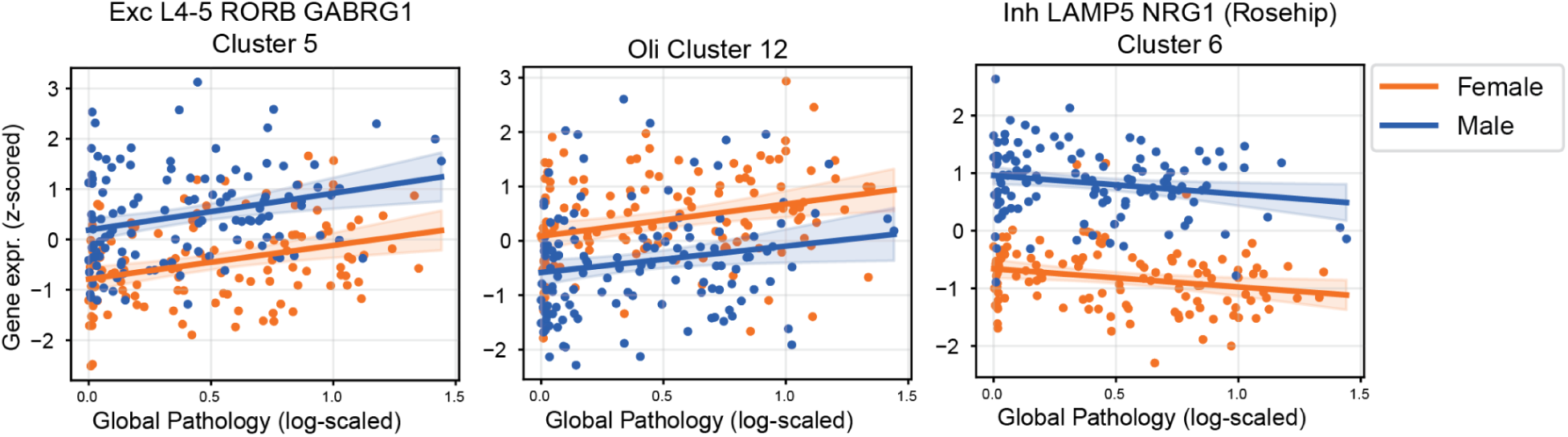
General additive models. (GAMs) showing gene modules that show statistically significant independent associations with both sex and global AD pathology. Orange indicates women and blue indicates men. Shaded regions represent 95% confidence intervals. Each point represents an individual donor.

## DISCUSSION

Alzheimer’s disease (AD) is increasingly understood as a multicellular disorder, in which amyloid and tau pathology accumulate alongside coordinated changes in various brain cells. Yet, it remains difficult to distinguish which transcriptional changes reflect causal disease-driving mechanisms, compensatory responses, or downstream consequences of neurodegeneration. Single-nucleus RNA sequencing makes it possible to reconstruct the natural history of these changes in human AD, moving toward models of how cellular and molecular responses emerge, interact, and change with pathology ^3,4,8,11^. These studies reveal widespread changes across diverse brain cell types, suggesting that AD-associated transcriptional variation is likely organised into complex, multicellular programs, rather than several populations changing separately. Modeling these programs will be essential for testing which cellular responses, if any, contribute causally to disease progression, how cells respond to drug treatments, and whether those changes mediate pathological or cognitive outcomes. These endeavors will not be possible without robust models of disease associated cellular changes that are reproducible across different cohorts.

We identified a robust network of transcriptomic modules, distributed across multiple cell types, that emerged in response to advancing AD pathology consistently across datasets. In contrast to some previous studies ^6,11,21^, the signature we identified was predominantly neuronal, with only a few modules stemming from glia. However, consistent with several previous studies identifying RORB-expressing excitatory neurons as selectively vulnerable in AD, including in entorhinal and neocortical excitatory populations affected by tau pathology ^4,7,22^, we too have detected AGMs from 3 different RORB-expressing subtypes, including RORB CUX2, RORB PLCH1 and RORB GABRG1. In particular, we found Exc L4-5 RORB GABRG1 neurons to contribute substantially more independent transcriptional modifications than other cell subtypes, suggesting these neuronal types have a particularly strong response to AD in the dlPFC. One of the positively-associated gene modules from this subtype was enriched for chemical synaptic transmission and G protein-coupled glutamate receptor signalling, and contained metabotropic glutamate receptor modulatory subunits *GRM3/4/8*, kainate receptor *GRIK2*, presynaptic genes *RIMS1* and *SYN3*, and several synaptic adhesion and guidance genes from *CDH*, *CNTN* and *CNTNAP* families. Together, these changes point to substantial synaptic dysregulation involving glutamatergic receptor signalling, synaptic remodelling, neurotransmitter release and neuronal excitability, all of which have been widely implicated in AD, including in neurons bearing neurofibrillary tangles ^22,23^. Similarly, previous studies have described an upregulated glutamatergic-neuron module in human AD enriched for voltage-gated calcium channels, GPCRs and increased predicted intracellular communication ^24^. However, RORB-associated vulnerability and transcriptional responses appear to vary by subtype and brain region. Mathys et al. identified a vulnerable RORB-expressing L5 AGBL1/GPC5 population in the entorhinal cortex, while Leng et al. found two vulnerable RORB-expressing EC subtypes and noted that the vulnerable RORB+ populations in the SFG were those most similar to their EC counterparts ^3,7^. In contrast, Dharshini et al. identified a distinct, relatively preserved RORB/CUX2/EYA4 layer 4 population with increased expression of genes involved in synaptic maintenance, plasticity and calcium homeostasis, which they interpreted as a compensatory response to hyperexcitability ^25^. Altered expression of *GRIK2*, *GRM5/7*, *NRXN1* and *CDH12* has also been linked to disturbed calcium signalling and firing homeostasis in human-induced glutamatergic neurons, with overlapping glutamate-receptor and channel alterations observed in glutamatergic neurons from human AD brains ^26^. Therefore, it is difficult to define whether mechanisms captured in RORB GARBG1 excitatory neurons represent compensatory maintenance of remaining connections, maladaptive excitatory signalling, or both.

This RORB GABRG1 signature was accompanied by transcriptomic changes in LAMP5 NRG1 (Rosehip) interneurons, which was another major source of transcriptomic variation observed in our analyses. LAMP5 NRG1 (Rosehip) interneurons had a module enriched in mitochondrial respiration and oxidative phosphorylation, driven by electron transport chain genes from *NDUF*, *UQCR*, and *COX* families, alongside superoxide-response and cell-death pathways involving *PRDX1*, *SOD1*, *GPX4*, *FTH1*, *CLU* and *HSP90AA1*. This module was negatively associated with AD pathology, suggesting reduced interneuron activity and stress-response capacity as pathology increases. Importantly, our CCC analysis predicted ligand-receptor interactions between RORB GABRG1 excitatory neuron and LAMP5 NRG1 (Rosehip) interneuron gene modules, including guidance, remodelling and neuromodulatory signals. Similar AD-associated excitatory-inhibitory communication changes involving synaptic signalling have recently been reported ^24^. An elevated synaptic excitation/inhibition (E/I) ratio has previously been observed in human AD parietal cortex across anatomical, electrophysiological and transcriptomic measurements, with the transcriptional and cellular shifts driven largely by reduced inhibitory components ^27^. LAMP5-expressing interneurons have been shown to suppress neuronal network hyperexcitation, while their loss contributes to AD-and tauopathy-related circuit dysfunction ^9^. The coupled RORB GABRG1 and LAMP5 NRG1 (Rosehip) signatures may therefore reflect a local E/I imbalance in which declining interneuron function permits or amplifies hyperexcitability-related changes in vulnerable RORB GABRG1 excitatory neurons.

Other neuronal and glial cell types, less central to this coordinated response, consistently showed strong enrichment for genes involved in axonogenesis and neuronal plasticity. A consistent observation across other neuronal and glial cells less prominent in this coordinated response, was a strong enrichment in transcription of genes involved in axonogenesis and neuronal plasticity. Mathys et al. also reported synapse-and cell projection related changes across multiple cell types ^4^, indicating broad pathology-associated changes in synaptic organisation and maintenance. Similar changes in axon-guidance and synaptic pathways have been observed in excitatory neurons during early amyloid pathology ^28^, while perturbed axonogenesis accompanied by impaired neurite outgrowth has been reported in iPSC-derived AD neurons ^29^. These recurring axonogenesis-related changes across several neuronal subtypes may reflect attempts to remodel or maintain neuronal connections as pathology progresses, but may also include maladaptive responses associated with impaired neurite growth. One interesting example of this in our dataset came from oligodendrocyte precursor cells (OPCs), which receive synaptic input from neurons, respond to neuronal activity, and contribute to circuit remodelling and activity-dependent myelination ^30–32^. Our CCC analysis revealed robust reciprocal communication between one prominent OPC module and several neuronal modules. This suggests that these OPCs may help maintain neuronal plasticity and circuit stability, with its pathology-associated decline contributing to the loss of neuronal support in AD. Similarly, Gabitto et al. reported early upregulation of OPC differentiation and remyelination factors, which they interpreted as a potentially compensatory response, followed by a decrease later ^8^. SMOC1, an OPC-expressed extracellular matrix protein, is also altered early with Aβ plaque accumulation and repeatedly emerges across CSF and brain proteomic studies of AD ^33–35^. In parallel, Aβ plaque-associated OPCs have been shown to adopt senescence-like inflammatory states and lose differentiation capacity ^36^, suggesting that the negative association of OPC C16 may reflect erosion of an OPC-mediated neuronal maintenance or plasticity state as pathology increases.

Microglia in AD are often associated with neuroinflammatory signatures ^37,38^. Whilst we didn’t detect any robust neuroinflammatory signatures in our results, we found two microglial gene modules that were negatively associated with AD and contained networks associated with axonogenesis, neuronal plasticity, and myelination. Previous literature has also shown that microglia are known to participate in synaptic remodelling and maintenance ^39,40^, and promote learning-dependent synapse formation through BDNF ^41^. It is worth noting that our use of cell-type specific background sets in our enrichment analyses may have attenuated our power to detect very common microglial signals, such as those related to neuroinflammation. Similarly, our null modeling approach seemed to disproportionately prune microglial CCC connections. We therefore cannot rule out the lack of expected neuroinflammation signals, due to our conservative approach resulting in type II error. Nevertheless, the negative association of synaptic maintenance microglial gene modules may reflect reduced homeostatic microglial support for neuronal processes along the pathology axis.

In all, increased AD pathology was associated with a strong transcriptomic response selectively in RORB GABRG excitatory neurons and LAMP5 inhibitory neurons, perhaps reflecting an E/I imbalance, and accompanied by reduced homeostatic functions in glial support cells, and increased synaptic reorganization in other neuronal types. Determining the biological role of this multicellular signature is inherently challenging. Observational transcriptomic data cannot establish whether it represents a compensatory response, a disease-driving mechanism, or a downstream consequence of accumulating pathology, and these possibilities are not mutually exclusive. Nevertheless, several features of our results provide clues to the biological significance of this cellular signature of AD. The signature appears organised, involving multiple different cell types and spanning across the pathology continuum. Together with the observed cell-cell communication network, this shows a reproducible tissue-level response that evolves throughout disease progression. This interpretation is consistent with recent transcriptomic, spatial, and proteomic studies showing that coordinated molecular networks emerge early in AD and continue to change across the pathological continuum ^8,10,11,33,42^. The signature also appears to contain both potentially adaptive and maladaptive components. Several positively associated AGMs were enriched for axonogenesis, neurite guidance, extracellular matrix remodelling and neuronal plasticity, while oligodendrocyte-associated modules furthermore showed enrichment for processes including ion homeostasis, intracellular signalling, lipid metabolism, extracellular remodelling and stress adaptation. These changes could represent attempts to preserve neuronal connectivity and tissue homeostasis in the face of accumulating pathology. Conversely, negatively associated AGMs suggest progressive loss of supportive cellular functions, including the decline of OPC modules enriched for axonogenesis and synaptic support, together with the reduction of LAMP5 rosehip interneuron modules enriched for mitochondrial and oxidative phosphorylation pathways. At the same time, multiple RORB GABRG1 excitatory neuron AGMs showed increased expression of synaptic signalling pathways, raising the possibility that selective loss of inhibitory restraint alongside enhanced excitatory signalling contributes to a gradual shift in cortical E/I balance, rather than simply widespread neuronal dysfunction. Taken together, these observations suggest that the signature reflects a structured change of the cortical cellular ecosystem, in which adaptive and maladaptive responses coexist throughout disease progression.

Finally, the pathology-related PLS axis was associated with clinical disease severity, while the mediation and moderation analyses indicated a more complex relationship between the multicellular signature, neuropathological burden and cognition. The signature did not mediate the effect of global AD pathology on cognition, arguing against a simple model in which pathology induces the multicellular response, which then drives cognitive decline. Instead, global AD pathology mediated the association between the signature and cognition, while the relationship between the signature and cognitive decline became stronger as pathology increased. This suggests that the signature may not be inherently detrimental, but that the cellular changes it captures become increasingly maladaptive as pathology accumulates. Similar transcriptomic studies have identified molecular states associated with both AD pathology and cognitive decline, while showing that their relationships with cognition are not fully accounted for by amyloid and tau burden alone ^4,43^. Rather than representing either successful compensation or inevitable degeneration, this coordinated multicellular signature may therefore capture how the brain reorganises as AD pathology accumulates, with different components contributing to tissue adaptation, cellular dysfunction and the clinical expression of disease.

One of the strengths of this study is the focus on optimizing data preprocessing in advance of data-analysis. Benchmarking studies have shown that parameter choices and workflow structure in snRNA and scRNA data contexts substantially affect downstream performance, and that optimal settings depend on the dataset, method, and analytical task, rather than being universal ^15,16,44–48^. Although much of this work has focused on clustering, dimensionality reduction, cell-type annotation, or tool comparison, the same principle is relevant for disease-focused modelling. This is also important for robustness, as single-cell studies can be vulnerable to false discoveries when replicate structure and technical variability are not handled carefully ^49,50^. In line with this, we did not treat default preprocessing settings as inherently best, but optimised parameters for the specific aim of identifying reproducible cell-subtype-specific gene modules that improved PLS model performance. Importantly, we did this in a manner that avoided data leakage and minimized overfitting. Thus, the selected parameter set should be interpreted as the best-performing configuration for this downstream task, and not as a generally superior preprocessing strategy for all snRNA-seq analyses. This optimisation improved not only apparent model performance but also generalisability. Concordant AGM loadings and successful projection of the ROSMAP-trained model into SEA-AD suggest that the workflow captured a reproducible biological structure rather than cohort-specific one. What was not necessarily expected was that the parameter set optimized in ROSMAP also performed substantially better than default settings in the SEA-AD dataset. While we cannot extrapolate whether this optimized parameter set is superior beyond the study of AD-associated changes in these two datasets, it does suggest that parameter optimization may generalize to additional datasets as long as the data and analytic goals remain similar.

The best-performing parameter sets converged on biologically interpretable features. Higher dispersion thresholds reduced the number of highly variable genes entering downstream analysis, suggesting that stricter feature selection helped limit noisy or weakly informative signals. In addition, higher gene-clustering ranges produced smaller and more specific AGMs, which may have helped separate AD-associated processes, such as stress responses, axonogenesis, and extracellular matrix remodelling, that would otherwise be collapsed into broader transcriptional aggregates. Subtype-specific metacells provided a compromise between robustness and biological resolution, sitting between noisy individual nuclei and coarse donor-level pseudobulking. This reduced dropout and sampling noise while retaining finer within-subtype variation ^51,52^. However, this robustness may come at the cost of sensitivity to rare or transient cell states, such as reactive microglial populations outnumbered by more abundant homeostatic microglia. Constructing metacells within predefined subtypes limited the risk of merging biologically distinct populations, although heterogeneous metacells can still distort downstream signals if not carefully tuned ^53^.

Sexual dimorphism in the prevalence of AD has been consistently reported and well described, with women more frequently affected than men ^54^. Women have also been reported to show faster cognitive decline and greater brain atrophy ^55,56^, more rapid tau accumulation in the presence of elevated amyloid burden ^57^, and differences in cognitive and neuropsychiatric presentation across the AD continuum ^55,56^. We were interested in characterizing a universal cellular response to AD, and so we initially included sex primarily as a covariate of no interest when defining AGMs. Despite this analytical choice, sex emerged as a strong and reproducible source of multicellular transcriptional variation. After the primary AD-related cellular response, the strongest source of covariance among AGMs related almost exclusively to sex, a finding that was replicated in the external SEA-AD cohort. While we detected no significant sex-by-pathology interactions among individual AGMs, several modules were independently associated with both sex and global AD pathology and showed sexually dimorphic baseline expression. Previous single-cell studies have primarily linked sex differences in AD to immune and glial transcriptional states, including alterations in microglial lysosomal activity ^58^, an increased abundance of disease-associated microglia in females ^19^, and astrocytic inflammatory responses ^59^. Our findings extend these observations by identifying sex-associated modules in oligodendrocytes and selected neuronal types, suggesting that sexual dimorphism may span a broad range of AD-relevant cellular processes. Some of these AGMs also contained sex-chromosome genes, suggesting that dimorphic expression may be influenced by differences in X and Y-linked gene expression as well as hormone-dependent regulation. However, these effects cannot be clearly disentangled using postmortem transcriptomic data alone.

One such example was oligodendrocytes cluster 12, which was positively associated with pathology and showed higher baseline expression in women. Although the module was not significantly enriched, several genes and a small subnetwork pointed to lipid and fatty-acid handling, and oxidative-stress responses. Its higher expression in women may reflect a more disease-associated or stress-responsive oligodendrocyte state at baseline, consistent with experimental evidence that sex-biased oligodendrocyte responses can emerge before myelin loss and vary with age and pathological context ^60^. Given the large number of poorly annotated transcripts in this AGM, it may also capture sex-related oligodendrocyte mechanisms that are relevant to AD but remain incompletely understood. Sexual dimorphism was also evident in the previously discussed LAMP5 NRG1 (Rosehip) response. Cluster 6 contained a subnetwork related to the unfolded-protein response, chaperone-mediated protein refolding, MAPK regulation and activity-dependent transcription, suggesting a stress-adaptive neuronal state ^61^. Its lower expression in women may therefore indicate reduced capacity to mount these responses, potentially increasing the susceptibility of LAMP5 NRG1 (Rosehip) interneurons to dysfunction as pathology accumulates. These observations raise the possibility that sex influences AD not only by modifying disease-associated responses, but also through inherent differences in the expression of gene networks that later participate in the disease process.

Several limitations should be considered when interpreting these findings. First, our analysis is based on postmortem single-nucleus transcriptomic data and therefore captures RNA abundance within surviving nuclei at a single terminal time point, rather than dynamic molecular changes in living tissue. Postmortem interval and other events surrounding death may further influence the measured expression profiles, despite statistical adjustment and quality control ^62^. Second, nuclear RNA represents only part of the cellular transcriptome and doesn’t reflect cytoplasmic or locally translated synaptic processes. Moreover, the most vulnerable cell populations may already have been depleted by the time of tissue collection, thus potentially biasing the observed signatures towards surviving cellular states. Our analyses were also limited to the dlPFC, a region affected relatively late in AD, which may limit detection of earlier disease-associated cellular changes. Our deliberately conservative discovery and validation framework prioritised robustness and replication over sensitivity. Although this reduced the likelihood of false-positive findings, it may also have excluded weaker, heterogeneous, or less well-powered disease-associated responses, particularly in rare cell populations. This may also partially explain why reproducible associations were concentrated among pathology measures, while APOE genotype and vascular measures showed few or none. This lack of robust associations likely reflects limited power, given the small number of E2 and E4 carriers, or weaker, more heterogeneous transcriptional effects, rather than a true absence of transcriptional associations. Biological interpretation was further limited by incomplete annotation resources, as many detected transcripts remain poorly characterised, while cell-cell communication analyses rely on ever evolving ligand-receptor databases that may still be missing relevant interactions. Since our snRNA-seq data lack spatial information, we cannot establish whether the predicted ligand-receptor pairs reflect direct interactions between cells in the tissue. Finally, generating sufficiently large donor-by-subtype groups for metacell and AGM construction required merging some related cellular states, preventing separate analysis of potentially distinct reactive populations, particularly among microglia.

In summary, we identified a reproducible multicellular transcriptional signature associated with AD pathology and replicated its major pathology-and sex-related axes in an independent cohort. The signature linked vulnerable neuronal populations with glial states involved in cellular stress, circuit regulation and neuronal support, suggesting that AD progression involves both adaptive and dysfunctional tissue-level responses. Future work should determine how specific this signature is to AD by examining aging and other neurodegenerative disease datasets, and assess how consistently the response is observed across different brain regions. Developing animal or cellular models that reproduce this multicellular response could help establish its temporal sequence, while analysis of tissue from patients treated with AD-targeting therapies may reveal whether modifying pathology alters the signature. Spatial and multi-omic approaches could be used to test the predicted cellular interactions and determine how the identified modules relate to protein-level changes, while experimental perturbation will be needed to establish which responses contribute to pathology and which reflect compensation. Overall, this framework provides a basis for studying AD as a coordinated multicellular process and for identifying cellular responses that may influence vulnerability, progression and clinical outcomes.

## Funding/Acknowledgements

This work was supported by the SciLifeLab & Wallenberg Data Driven Life Science Program (grant: KAW 2020.0239). G.V. also received travel grants from the Strategic Research Area MultiPark (Multidisciplinary Research on Neurodegenerative Diseases) at Lund University and the Alzheimer’s Association to present this work. The results published here are in whole or in part based on data obtained from the AD Knowledge Portal (NIH-NIA, U24AG061340; https://adknowledgeportal.org/). Study data were generated from postmortem brain tissue provided by the Religious Orders Study and Rush Memory and Aging Project (ROSMAP) cohort at Rush Alzheimer’s Disease Center, Rush University Medical Center, Chicago. This work was supported in part by the Cure Alzheimer’s Fund, NIH grants AG058002, AG062377, NS110453, NS115064, AG062335, AG074003, NS127187, MH119509, HG008155 (M.K.), RF1AG062377, RF1 AG054321, RO1 AG054012 (L.-H.T.) and the NIH training grant GM087237 (to C.A.B.). ROSMAP is supported by P30AG10161, P30AG72975, R01AG15819, R01AG17917. U01AG46152, U01AG61356. The ROSMAP single-nucleus RNA-sequencing data used in this study were obtained under Synapse accession syn52293417. The Seattle Alzheimer’s Disease Brain Cell Atlas (SEA-AD) consortium is supported by the National Institutes on Aging (NIA) grant U19AG060909. Study data were generated from postmortem brain tissue donated to the University of Washington BioRepository and Integrated Neuropathology (BRaIN) laboratory and Precision Neuropathology Core, which is supported by the UW Alzheimer’s Disease Research Center (NIA P30AG066509, previously P50AG005136), the Adult Changes in Thought (ACT) Study (NIA U19AG066567). ACT Data collection for this work was additionally supported, in part, by prior funding from NIA (U01AG006781), and the Nancy and Buster Alvord Endowment. We thank the participants of the ADRC and the ACT study for the data they have provided and the many ADRC and ACT investigators and staff who steward that data. You can learn more about UW ADRC and ACT. We thank AMP-AD, Sage Bionetworks and the Open Data Registry on AWS for hosting various datasets from this study.

## Conflicts of Interest

The authors declare no conflicts of interest.

## Data Availability

The ROSMAP single-nuclei RNA-seq dlPFC data was obtained from the AD Knowledge Portal on Synapse (accession number: syn52293433), under the required controlled-access data use agreement.SEA-AD single-nucleus RNA-sequencing data were obtained from the publicly available SEA-AD Single Cell Profiling dataset through the AWS Open Data Registry in (sea-ad-single-cell-profiling) as AnnData objects (h5ad files).

## Code Availability

All original code has been deposited to GitHub and is available for viewing (https://github.com/DeMONLab-BioFINDER/MC_characterisation).

## Author Contributions

GV and JWV conceptualized the study. GV, LA, NMC, CG, and JWV contributed to analytical design. GV, EV, and YX involved in data processing and curation. GV and EV conducted data analysis. JWV supervised the study. GV and JWV wrote the manuscript. All authors interpreted the data and contributed substantively to revising the manuscript.

## METHODS

### ROSMAP dataset and donors

We analyzed single-nucleus RNA-seq (snRNA-seq) data (∼2.3 million nuclei) from the dorsolateral prefrontal cortex (DLPFC) of 427 post-mortem donors from the Religious Orders Study and Memory and Aging Project (ROSMAP) cohort ^4^. ROSMAP is a community-based convenience sample: participants in the Religious Orders Study were Catholic nuns, priests, and brothers from across the United States, while those in the Memory and Aging Project were older adults recruited from retirement communities and subsidized housing in the Chicago area. All participants were free of dementia at enrollment, underwent annual clinical evaluations, and consented to brain donation at death under protocols approved by the institutional review board of Rush University Medical Center. Participants also signed an Anatomic Gift Act, and a repository consent to allow their data to be repurposed.

Donors were older adults with an average age at death of ∼89 years, and sex was nearly evenly distributed (215 females, 212 males). Clinical diagnoses included cognitively unimpaired (CU, *n* = 146), mild cognitive impairment (MCI, *n* = 110), Alzheimer’s disease (AD, *n* = 160), and other dementia cases (*n* = 11). For the purpose of analysis, AD and other dementia cases were combined into a single category.

Extensive metadata were available for all donors, including demographic information (age at death, sex), APOE genotype, and a range of neuropathological measures: amyloid load and neurofibrillary tangle density, as well as Braak stage, CERAD neuritic plaque score, cerebral amyloid angiopathy, arteriolosclerosis, cerebral atherosclerosis, chronic microinfarcts, and global AD pathology burden, which is a quantitative composite measure derived from counts of neuritic plaques, diffuse plaques, and neurofibrillary tangles across multiple brain regions ^63^.

### Data acquisition

As reported by Mathys et al. (2023), nuclei were isolated from frozen post-mortem DLPFC tissue prepared using droplet-based 10x Genomics Chromium Single Cell 3’ Reagent Kits v3 according to the manufacturer’s protocol. The snRNA-seq libraries were sequenced using NextSeq 500/550 High Output v2 kits (150 cycles) or NovaSeq 6000 S2 Reagent Kits.

Gene counts were acquired by aligning reads using Cell Ranger v3.0.2 against the GRCh38 genome. Following quality control and doublet removal cell identities were assigned based on previously published snRNA-seq data (Allen Institute’s cell types database; https://portal.brain-map.org/atlases-and-data/rnaseq/human-multiple-cortical-areas-smart-seq). For the present study, raw gene count matrices and the corresponding cell type and subtype annotations were accessed through the AD Knowledge Portal on Synapse (https://www.synapse.org/#!Synapse:syn52293417) and used as input to the analysis pipeline.

### SEA-AD cohort replication

For external replication, we used snRNA-seq data from Seattle Alzheimer’s Disease Brain Cell Atlas (SEA-AD) ^8^. The available dataset contains approximately 1.6 million nuclei from the prefrontal cortex of 80 post-mortem donors, including 49 females and 31 males, with a mean age at death of ∼88 years. Extensive donor-level metadata were available, including measures of AD neuropathology such as Braak stage, APOE genotype, and CERAD score, as well as comorbid neuropathologies including atherosclerosis, arteriosclerosis, and microinfarcts. Detailed donor inclusion and exclusion criteria are described in Gabitto et al. (2024). Informed consent was obtained according to protocols approved by the University of Washington and Kaiser Permanente Washington Health Research Institute Institutional Review Boards ^8^.

### Pipeline overview

We developed a multi-step computational pipeline to identify robust gene expression modules associated with Alzheimer’s disease pathology from single-nucleus RNA-seq data (Fig. 1A). Before running the main pipeline, a preliminary preprocessing and clustering step was used to consolidate low-abundance cell subtypes that contained too few nuclei to support stable downstream preprocessing. Following subtype consolidation, raw count matrices were reprocessed separately for each major cell type using the parameter settings evaluated in the optimization framework. Then the data was grouped by cell subtypes and donors, and transcriptionally similar nuclei were aggregated into metacells to reduce single-cell sparsity and generate more stable expression profiles. Donors were then split into three equal parts: train, validate and test sets, and gene co-expression modules were constructed within each subtype and tested for associations with disease-related phenotypes using linear mixed effects models. Modules that replicated across training and validation datasets were retained as AD-associated gene modules for downstream multivariate Partial Least Squares (PLS) and other downstream analyses. These steps are outlined in detail below.

### Data train, validation, test split

Dataset donors were assigned into three equal sets, stratified by sex and dementia diagnosis, with all nuclei and derived metacells from the same donor assigned to the same dataset. Training and validation sets were used to identify robust AD-associated gene modules, and the test set was used for PLS evaluation.

### Cell-subtype consolidation

Before running the main analysis pipeline, we performed a preliminary preprocessing step to identify low-abundance subtypes requiring consolidation. ROSMAP single-nucleus RNA-seq count matrices were processed at each major cell type level using Scanpy v1.10.1 ^64^. Nuclei with fewer than 100 detected genes were removed, and genes detected in fewer than three nuclei were excluded. Counts were then library-size normalized to 10,000 counts per nucleus and log-transformed. Highly variable genes were identified separately within each major cell type initially using minimum and maximum mean-expression thresholds of 0.0125 and 3.0, and a minimum dispersion threshold of 0.5. We then calculated cell subtype-averaged HVG expression profiles and performed hierarchical clustering using Ward linkage and Euclidean distance. Cell subtypes represented by fewer than 20,000 nuclei across all donors were merged with their nearest transcriptional neighbor. The complete mapping from original annotations to merged subtypes can be found in Supplementary Table 7. This preliminary preprocessing step was used only to determine the final subtype groupings. The normalized expression values and highly variable genes generated during this step were not carried forward into the main analysis pipeline.

### Cell type harmonization between SEA-AD and ROSMAP

Cell subtype naming conventions differed between the ROSMAP and SEA-AD datasets. However, because the underlying transcriptional subtype structure showed high concordance across cohorts, SEA-AD subtype labels were harmonized to match the ROSMAP nomenclature before downstream replication analyses. Cell subtype matching was performed separately within each major cell type. For both datasets, nuclei with fewer than 100 detected genes were removed, and genes detected in fewer than three nuclei were removed. Genes shared between ROSMAP and SEA-AD were then retained. Expression values were library-size normalized to a target sum of 10,000 counts per nucleus and log-transformed. For each dataset, mean gene expression was then calculated for each subtype using the shared gene set. Subtype-level expression profiles were standardized by z-scoring, and pairwise correlations were computed between ROSMAP and SEA-AD subtype profiles. The resulting correlation matrices were visualized as heatmaps and manually inspected to identify the best-matching SEA-AD subtype for each ROSMAP subtype. These matches were then used to rename SEA-AD subtypes according to the ROSMAP annotation scheme (Supplementary Fig. 7; Supplementary Table 7).

### Data processing and quality control

Following subtype consolidation, the raw count matrices were reprocessed from the beginning separately for each major cell type, since downstream analyses compared gene networks within specific cell subtypes rather than across the entire dataset. Nuclei and lowly detected genes were removed, after which counts were library-size normalized to 10,000 counts per nucleus and log-transformed. Highly variable genes (HVGs) were identified using the function *scanpy.pp.highly_variable_genes()* at the major cell type level. HVG selection served as a feature selection step, restricting analyses to the most informative genes while reducing noise from invariant ones. Additionally, identifying HVGs at the cell type (rather than subtype) level ensured that gene lists were specific to each major lineage but remained comparable across its constituent subtypes. Parameter optimization for HVG selection is described below, in section *Parameter optimization*).

### Metacell generation

Because snRNA-seq data is known to be noisy and highly sparse, largely due to stochastic dropout ^65,66^, we aggregated cells into metacells to reduce this noise and sparsity (Fig. 1A). This pooling strategy differs from pseudobulking, which groups all cells of a subtype within a donor into one profile. Metacells instead group a predefined number of similar cells, preserving within-donor cell heterogeneity while still producing more stable expression profiles that are better suited for downstream analyses ^52^.

To achieve this, metacells were constructed within each donor and cell subtype group using a k-nearest neighbour (kNN) graph. The parameter *k* (number of neighbors) controlled granularity, with smaller values retaining fine-grained variation and vice versa. *k* was defined via parameter optimization, described below. For each donor-by-subtype group, metacells were generated iteratively. At each iteration, PCA was performed on the remaining cells to identify a seed cell, defined as the cell with the largest absolute PC1 score. The seed cell and its nearest neighbors, for a total of k cells, were grouped into one metacell and then removed from the remaining cell pool. This procedure was repeated until no further metacells could be formed. After metacell clustering, each metacell was assigned to the training, validation, or test set according to the donor-level split described earlier. Metacell sparsity was calculated as the proportion of genes with zero detected expression, and metacells exceeding the predefined sparsity level (an optimised parameter) were removed.

### Gene co-expression clustering

Individual gene expression is noisy and expression changes rarely happen in isolation. Therefore, after clustering the cells into metacells, we then clustered the genes into gene co-expression modules via *k*-means to derive co-expression networks that may reflect independent intracellular processes, pathways or components (Fig. 1A). The optimal number of gene co-expression modules was determined by scanning across a defined range of *k* values (an optimized parameter), calculating silhouette scores for each solution, and selecting the peak with the highest score. Module assignments defined in the training set were then propagated to the validation and test sets.

### Association testing with linear mixed-effects regression (LMER)

Gene clustering across all subtypes produced a large number of modules, many of which were unlikely to be relevant to AD. To reduce dimensionality and focus on biologically meaningful signals, we tested each module for association with AD-associated phenotypes using linear mixed-effects regression (LMER), implemented in the lme4 and lmerTest R packages (Fig. 1A). The model included phenotype, sex, and sparsity as fixed effects and donor as a random effect:

***Gene co-expression module ∼ phenotype + sex + sparsity + (1|donor)***

Phenotypes included APOE E2 and E4 allele status, cognitive diagnosis, age at death, Braak stage, CERAD score, global AD pathology (gpath), NIA-Reagan score, amyloid load, neurofibrillary tangles, arteriolosclerosis, cerebral amyloid angiopathy (CAA), presence of microinfarcts, and cerebral atherosclerosis. Gene expression modules significantly associated with any phenotypes after FDR correction (FDR <0.05) in the training set were taken forward to the validation set, where replication was defined as passing a nominal p-value < 0.05. Only modules replicating in both datasets were retained as AD-associated gene modules (AGMs).

### Partial least squares (PLS) regression

Rather than investigating AGMs individually, we applied partial least squares (PLS) regression to jointly analyze all AGMs and AD phenotypes using a multivariate approach (Fig. 1). PLS derives latent components that maximize the covariance between two data matrices, allowing us to identify coordinated patterns of AGMs that covary with multiple disease traits simultaneously. In our case, we sought to estimate covariance between AGMs (X) and AD-related phenotypes (Y).

AGMs (PLS X) from all cell types were combined across the training and validation sets. For each AGM, metacell-level expression values were averaged per donor to generate a donor-level AGM expression matrix. Donors and AGMs with more than 30% missing values were excluded. Remaining missing values were imputed using k-nearest-neighbor imputation (*k* = 5), and AGM expression values were scaled using MinMax scaling. Phenotype data were processed using the same imputation and scaling approach. Amyloid, tangle, and global AD pathology scores were log-transformed before scaling.

The phenotype data (PLS Y) was restricted to a focused panel of age-and AD-relevant variables, including age at death, global AD pathology, amyloid load, neurofibrillary tangles, TDP-43 stages, arteriolosclerosis, cerebral amyloid angiopathy (CAA), cerebral atherosclerosis, chronic microinfarcts, APOE E2 and E4 allele status, and the final consensus cognitive diagnosis at the time of death (CU, MCI, or dementia).

To determine the optimal number of components in PLS model selection, models with one to ten components were trained and evaluated using five-fold cross-validation, repeated ten times. Adjusted cross-validated R2 and mean absolute error (MAE) were computed, and the model with the highest adjusted cross-validated R2 was selected. Statistical significance of the PLS components was assessed using permutation testing with 1,000 iterations. Stability of PLS component loadings was assessed using bootstrap resampling with replacement across 10,000 iterations.

### Parameter optimization

Because upstream preprocessing choices influence the AGM features used in the final PLS model, and there is no single universally optimal setting for these steps in single-nucleus RNA-seq data, we systematically tested several pipeline parameters and ranked parameter sets by downstream (cross-validated) PLS performance. Pipeline parameters were varied across HVG selection, metacell construction, sparsity filtering, and gene-module clustering. For HVG selection, several dispersion thresholds and minimum and maximum expression cutoffs were tested. For metacell construction, different *k* values and clustering resolutions were tested, including pseudobulking as a non-metacell comparison. We also varied the threshold used to exclude sparse metacells. For gene-module clustering, multiple *k* ranges were assessed to determine the optimal number of modules (Table 1). The full parameter grid was run in parallel on a high-performance computing cluster, testing 432 parameter combinations in total. Parameter sets were ranked by mean repeated adjusted cross-validated PLS R² within the merged training and validation set. The independent test set was withheld from parameter tuning and used only for final model assessment.

### Independent test-set evaluation

For final model evaluation, the PLS model was fitted on the merged training-validation dataset using the selected number of components. The independent test set was processed using the same selected AGMs and transformed using the imputation and scaling parameters fitted on the merged training-validation data. The trained PLS model was then applied to the transformed test set to evaluate generalizability.

### Reconstruction of ROSMAP-derived AGMs in SEA-AD

After subtype harmonization, the SEA-AD snRNA-seq dataset was processed using the default and optimized pipeline parameters selected in ROSMAP. Because SEA-AD was used as an external replication cohort, gene modules were not independently rediscovered in SEA-AD. Instead, the AD-associated gene modules (AGMs) previously defined in ROSMAP were reconstructed in SEA-AD using the same module gene assignments. This allowed us to test whether the transcriptional programs identified in ROSMAP could be recovered in an independent cohort.

### De novo PLS analysis in SEA-AD

To assess whether similar multivariate transcriptomic–phenotypic structure could be identified directly within SEA-AD, we trained a de novo partial least squares (PLS) regression model using SEA-AD donor-level AGM expression profiles as predictor variables. The phenotype matrix included age at death, sex, overall AD neuropathologic change, Braak stage, Thal phase, and CERAD score. Donor-level AGM expression values were derived as described for ROSMAP by averaging AGM expression across metacells belonging to the same donor. Model preprocessing, component selection, and evaluation followed the same procedure used for the ROSMAP PLS analysis.

### Projection of SEA-AD donors into the ROSMAP-derived PLS space

To directly assess the replicability of the ROSMAP-derived transcriptional structure, the fitted ROSMAP PLS model was applied to the SEA-AD donor-level AGM matrix using the *PLSRegression.transform()* function. This projects new observations into the latent space defined by the original ROSMAP model without refitting the model in SEA-AD.

Before projection, the SEA-AD predictor matrix was aligned to the ROSMAP AGM feature matrix to ensure that module columns matched the features used to train the original ROSMAP model. Missing values in the SEA-AD matrix were imputed using the gene imputer fitted to the ROSMAP training data, after which the matrix was transformed using the MinMax scaler fitted to the ROSMAP training data. Neither the imputer nor the scaler was refitted using SEA-AD data. One ROSMAP-derived module, Exc NRGN Cluster 15, was not present in the SEA-AD reconstructed AGM matrix. To preserve feature alignment, a placeholder column was added for this module. Because MinMax scaling was used during preprocessing, all values in this placeholder column were assigned a value of 0.5, corresponding to the midpoint of the scaled feature range. The resulting SEA-AD matrix was then transformed using the fitted ROSMAP PLS model to derive SEA-AD PLS X scores. These projected scores were used to evaluate whether the donor-level transcriptional axis identified in ROSMAP generalized to the independent SEA-AD cohort.

### Pseudoprogression analysis

To capture the trajectories of transcriptional changes across disease progression, we treated global AD pathology burden as a cross-sectional proxy for disease progression and fit each AGM as a smooth function of log-transformed global AD pathology using generalized additive models (GAMs) with the pyGAM package v0.12.0 ^67^. AGM expression values were z-scored to put them on a comparable scale, and models were fitted with penalized spline terms (splines = 5). P-values for the global AD pathology term were corrected across AGMs using false discovery rate (FDR) correction. To compare dynamic patterns across AGMs, fitted curves were predicted over a common global AD pathology grid and independently z-scored to remove differences in baseline expression and response amplitude, thereby emphasizing trajectory shape. Similarity between the standardized curves was quantified using cosine similarity, and AGMs were grouped into trajectory classes using spectral clustering. Because these data are cross-sectional rather than longitudinal, we refer to these curves as pseudoprogression trajectories.

### Sex-stratified modeling

To characterize how these transcriptional trajectories differ between men and women, we extended the GAM model by adding a sex term and an interaction term between globa lAD pathology burden and sex. For each AGM, we applied false discovery rate (FDR) correction for the p-values to identify modules with sex-global AD pathology interactions and independent associations with each variable.

### Cell-cell communication analysis

To infer putative intercellular signaling involving AGMs, we used the curated interaction annotations from the CellPhoneDB v5.0.0 database (accessed via: https://github.com/ventolab/CellphoneDB-data) ^68^. We did not use the CellPhoneDB analysis software; instead, we implemented custom code to identify CellPhoneDB-supported ligand-receptor, adhesion, and gap-junction interactions between genes contained in AGMs and their corresponding cell subtypes.

We performed two complementary permutation analyses to assess whether the observed AGM interaction network showed greater connectivity than expected by chance. For both analyses, we generated 10,000 null module sets in which each null module was matched to an observed AGM by module size and sampled from the same subtype-specific highly variable gene background. First, we tested overall network connectivity by counting the number of edges formed between modules in each null network and comparing this null distribution with the number of edges observed in the AGM network. In this analysis, edges were treated as binary: two AGMs were considered connected if at least one CellPhoneDB-supported interaction was identified between them, regardless of how many individual ligand-receptor, adhesion, or gap-junction interactions supported that connection. The observed AGM network was considered more connected than expected when its number of edges exceeded the upper bound of the 95% null interval. Second, we assessed edge strength by comparing the observed weight of each AGM-AGM connection, defined by the number of interactions, with the corresponding null distribution generated from matched null module pairs. Here we considered the edge significant when its weight exceeded the null 95% interval. The same framework was also applied at the parent cell-subtype level by aggregating module-level interactions according to their assigned subtypes, allowing us to identify subtype-subtype connections with greater interaction strength than expected by chance. These analyses were used to prioritize putative sender-receiver relationships associated with AD-linked transcriptional programs.

### Gene set annotation analyses

Gene set annotation analyses were performed to assign biological interpretations to the AGMs. First, each AGM was tested for functional and pathway enrichment using over-representation analysis (ORA). Analysis was performed using *gseapy.enrichr()* function from the Python gseapy v1.1.13 package ^69^. For each AGM, the gene input list consisted of genes assigned to that AGM and the background gene set was defined as the highly variable genes from the corresponding cell type from which the AGM was derived. For the enrichment, we queried KEGG and Gene Ontology Biological Process databases, and the terms were considered significantly enriched if the adjusted p-value was < 0.05 (FDR correction).

Additionally, since the AGMs were derived using a purely data driven approach, we additionally used STRING DB to see whether genes within each AGM formed smaller functionally connected subgroups. Networks were queried using the functional network type with a required interaction confidence score of 400. The STRING evidence channels retained for network construction were co-expression, experimental evidence, and text mining, while neighbourhood, fusion, co-occurrence, and database evidence channels were excluded. The resulting STRING edge tables contained pairwise gene associations and channel-specific evidence scores. Using NetworkX, we then grouped genes connected by STRING edges into groups, and using STRING gene set description API, the group functional descriptions were annotated based on STRING internal enrichment. For STRING-connected groups that remained unannotated, we performed an additional ORA analysis, using the same gseapy.enrichr() approach as described above.

Finally, AGM or STRING-connected subgroups that remained without statistically significant enrichment were assessed descriptively by looking at the gene composition. Known functional gene groups were used to provide qualitative labels where possible. AGMs without clear enrichment or interpretable gene-level themes were left unannotated. These approaches were considered to represent three decreasing levels of evidence: AGM-level enrichment, enrichment within STRING-connected subgroups, and descriptive interpretation based on gene composition.

### Moderation and Mediation analyses

Exploratory mediation and moderation analyses were performed to examine relationships among donor-level PLS component 1 (PLS C1) gene scores, global AD pathology burden, and retrospective longitudinal cognitive decline before death. For cognitive outcomes, we used estimated person-specific slopes for working memory, episodic memory, semantic memory, perceptual speed, perceptual orientation, and global cognition, as previously described ^70^. These slope measures represent the rate of change in cognition over time, and were previously adjusted for age at baseline, sex, and years of education. The underlying cognitive measures were composite scores derived from a battery of 19 neuropsychological tests. The tests contributing to each cognitive measure are provided in Supplementary Table 8.

Mediation analyses were performed using Pingouin v0.6.0 ^71^. Two complementary mediation models were tested for each cognitive domain. In the first model, global AD pathology was specified as the predictor, PLSX1 gene score as the mediator, and cognitive slope as the outcome. In the second model, PLSX1 gene score was specified as the predictor, global AD pathology as the mediator, and cognitive slope as the outcome.

Moderation analyses tested whether the association between PLSX1 scores and cognitive slopes varied at different levels of global AD pathology burden. Linear models were fitted with an interaction term between PLSX1 and global AD pathology, adjusting for age at death and sex. For visualization purposes, donors were stratified into low, medium, and high global AD pathology groups.

### Linear regression associations with regional pathology measures

To determine whether the DLPFC-derived PLSX1 gene score was associated specifically with pathology in the prefrontal cortex or reflected a broader brain-wide neurodegenerative process, we examined its associations with regional pathology measures. Here we’ve used linear regression models to associate PLSX1 gene scores with pathology whilst covarying for age at death and sex. Regional tau-related measures included neurofibrillary tangle burden and tau tangle measures across multiple brain regions: the angular gyrus, entorhinal cortex, hippocampus, midfrontal cortex, midtemporal cortex, calcarine cortex, anterior cingulate cortex, inferior temporal cortex, and superior frontal gyrus. Amyloid-related measures included amyloid burden, neuritic plaques, and diffuse plaques measured across the same available brain regions.

## Supplementary data

**Supplementary Figure 1.**
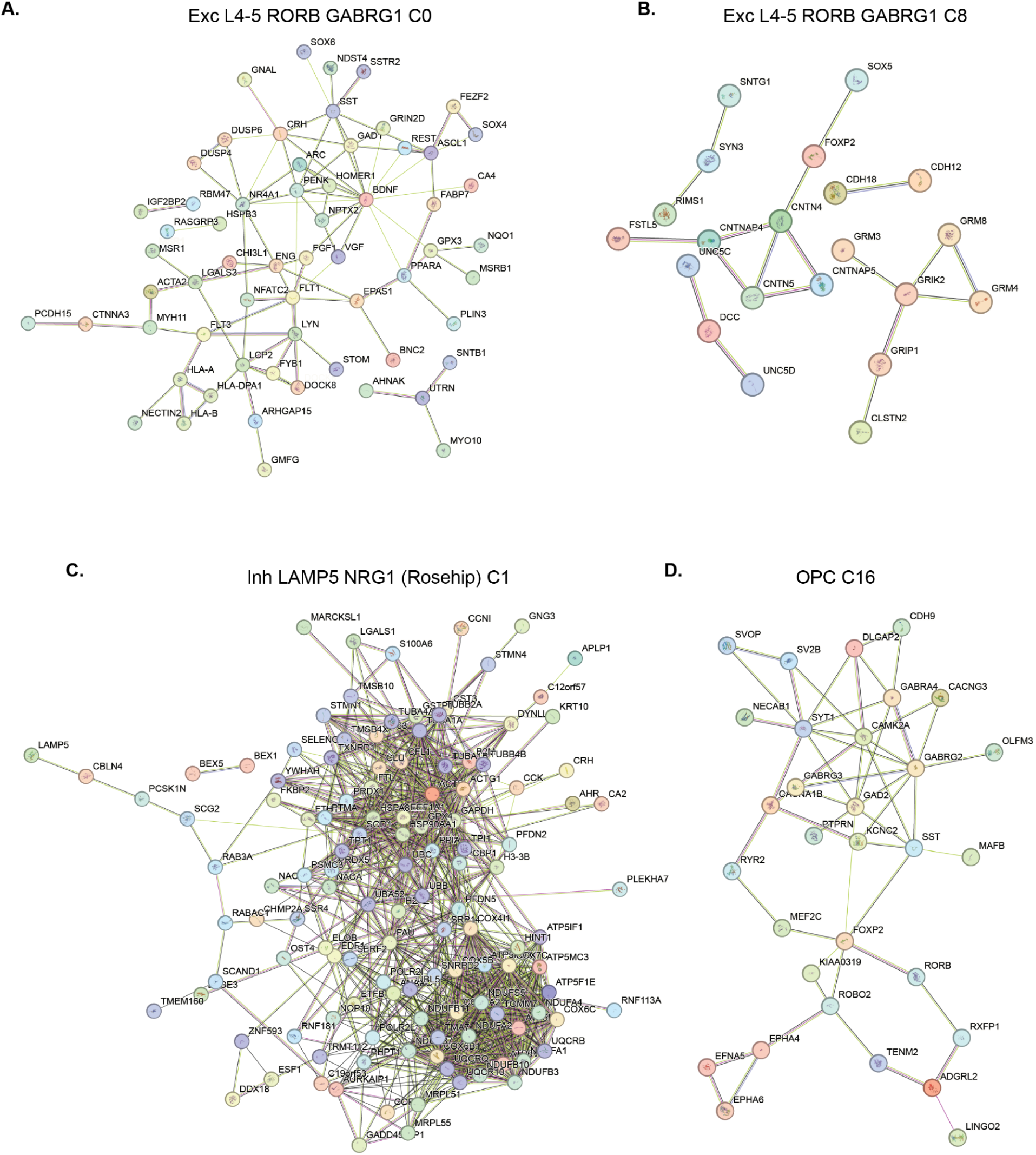
Representative STRING protein association subnetworks from selected AGMs. STRING functional association networks for **(A)** Exc L4-5 RORB GABRG1 C0, **(B)** Exc L4-5 RORB GABRG1 C8, **(C)** Inh LAMP5 NRG1 (Rosehip) C1 and **(D)** OPC C16. Networks were generated using a minimum STRING interaction score of 0.4 and evidence from co-expression, experimental data and text mining. Nodes represent proteins encoded by genes within each AGM; disconnected proteins are hidden. Edge colours indicate the supporting evidence source: black - co-expression; purple - experimental evidence; yellow - text mining.

**Supplementary Figure 2.**
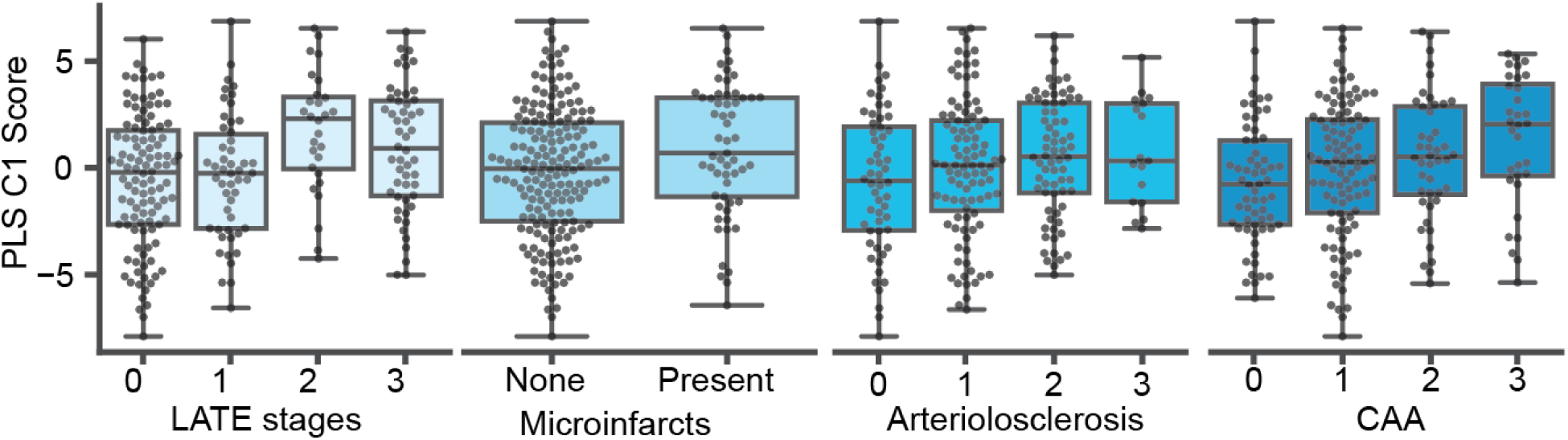
ROSMAP PLS Component 1 scores against associated pathology phenotypes.

**Supplementary Figure 3.**
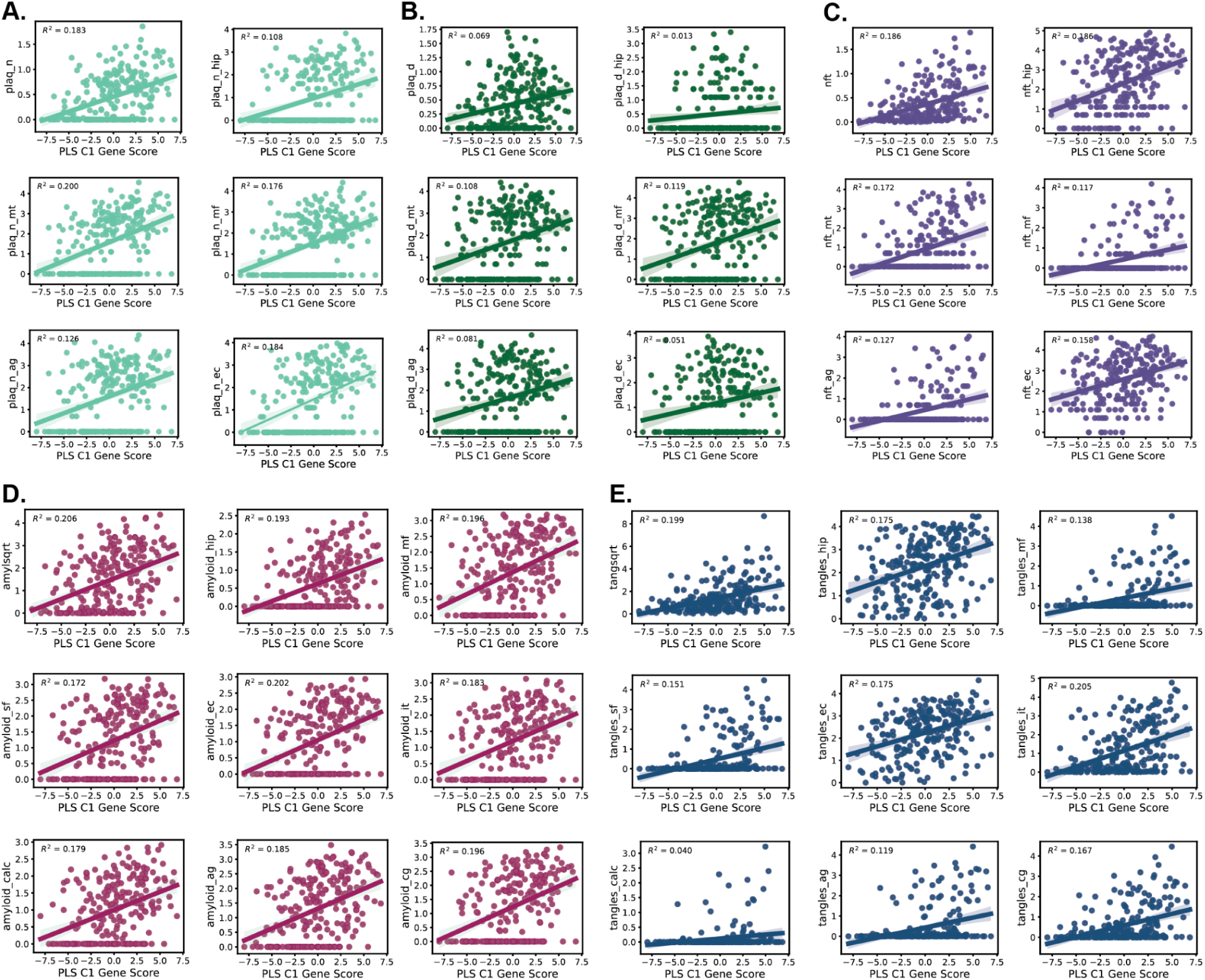
Associations between PLS-C1 gene scores and regional neuropathology measures. **(A)** neuritic plaques, **(B)** diffuse plaques, **(C)** neurofibrillary tangles (nft) **(D)** amyloid, and **(E)** tangles.

**Supplementary Figure 4.**
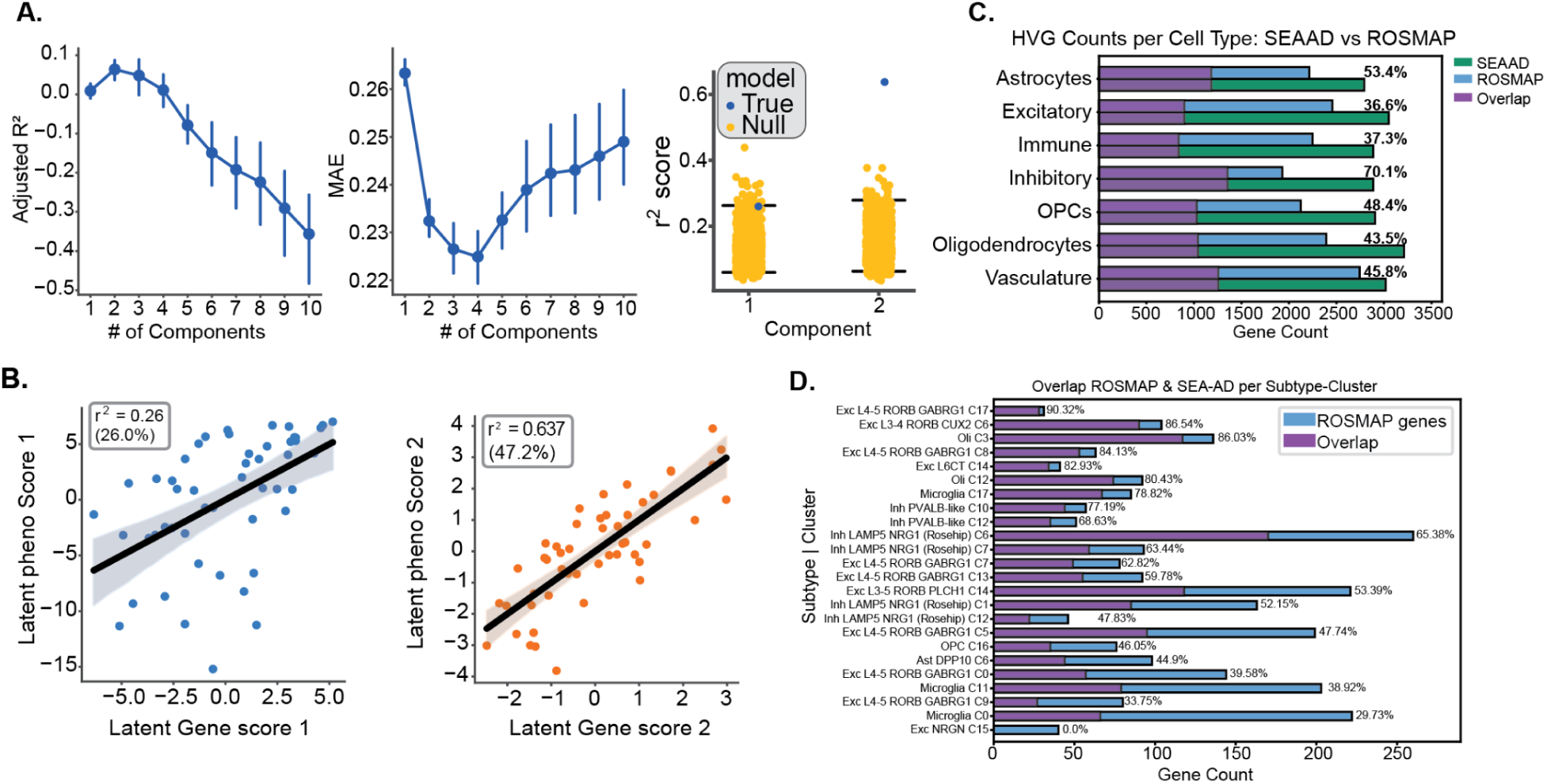
ROSMAP PLS model replication in external SEA-AD cohort. (A) SEA-AD PLS model performance. Left: Adjusted CV R2 scores across different numbers of components. Middle: mean absolute error (MAE) across different numbers of components. Right: showing the observed model R2 (blue) compared to null distributions (orange) for the selected model. **(B) Scatterplots of latent gene scores against latent phenotype scores** for PLS components 1 and 2, show the shared covariance captured by each component. **(C)** Overlap of highly variable genes between ROSMAP and SEA-AD within each major cell type. **(D) Reconstruction of ROSMAP AGMs in SEA-AD**, showing the total number of genes in each ROSMAP AGM and the subset also identified among the corresponding SEA-AD highly variable genes.

**Supplementary Figure 5.**
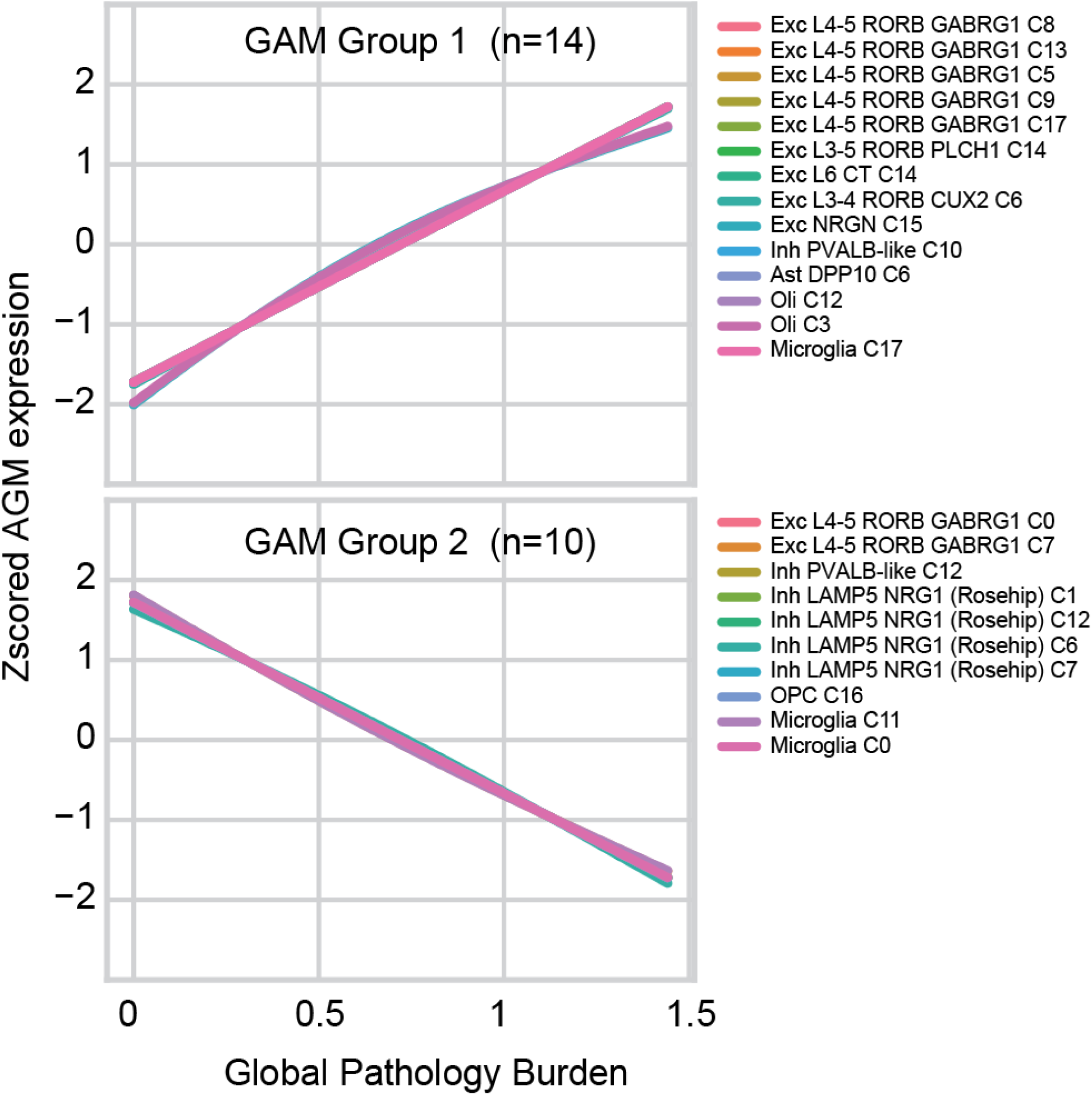
AGM pseudoprogression along the log-transformed global AD pathology burden, AGMs grouped by spectral clustering.

**Supplementary Figure 6.**
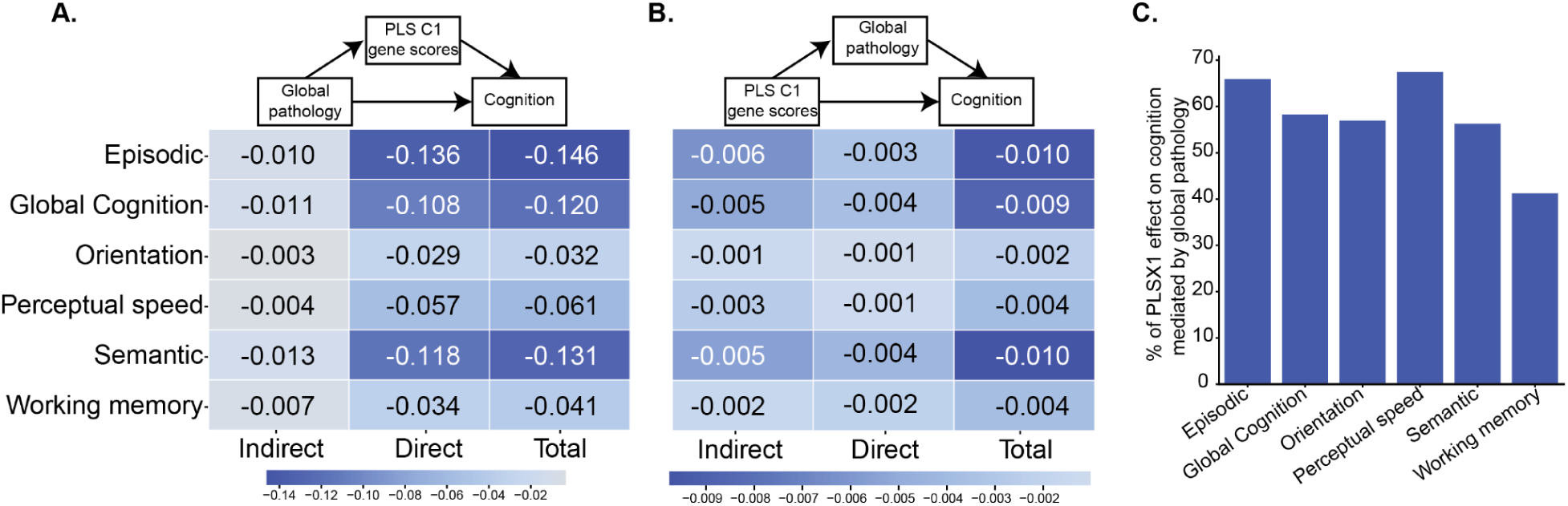
Mediation analyses linking global AD pathology, PLS-C1 gene scores, and longitudinal cognitive decline. **(A)** Mediation model testing whether PLS-C1 gene scores mediate the association between global AD pathology and cognitive decline. **(B)** Mediation model testing whether global AD pathology mediates the association between PLS-C1 gene scores and cognitive decline. Columns show indirect, direct, and total effects for each cognitive outcome. **(C)** Percentage of PLSX1 effect on cognition variables mediated by global AD pathology burden.

**Supplementary Figure 7.**
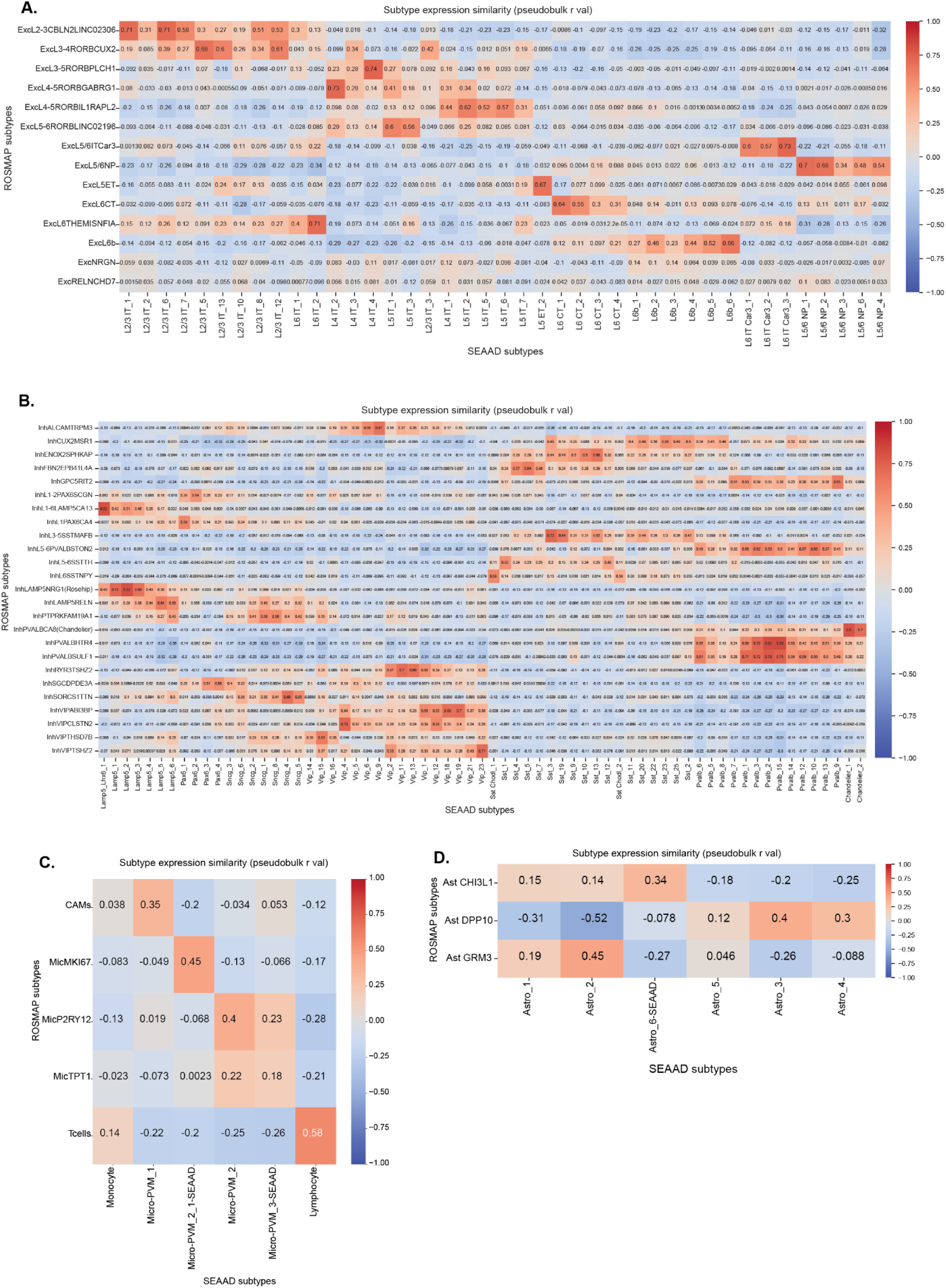
Cell subtype correlations between ROSMAP and SEA-AD datasets. **(A)** excitatory neurons, **(B)** inhibitory neurons, **(C)** microglia, and **(D)** astrocytes.

## Notes

### Competing Interest Statement

The authors have declared no competing interest.

## Bibliography

1. Brase, L., You, S.-F., D’Oliveira Albanus, R., Del-Aguila, J.L., Dai, Y., Novotny, B.C., Soriano-Tarraga, C., Dykstra, T., Fernandez, M.V., Budde, J.P., et al. (2023). Single-nucleus RNA-sequencing of autosomal dominant Alzheimer disease and risk variant carriers. Nat Commun 14, 2314.

2. Consens, M.E., Chen, Y., Menon, V., Wang, Y., Schneider, J.A., De Jager, P.L., Bennett, D.A., Tripathy, S.J., and Felsky, D. (2022). Bulk and Single-Nucleus Transcriptomics Highlight Intra-Telencephalic and Somatostatin Neurons in Alzheimer’s Disease. Front Mol Neurosci 15, 903175.

3. Mathys, H., Boix, C.A., Akay, L.A., Xia, Z., Davila-Velderrain, J., Ng, A.P., Jiang, X., Abdelhady, G., Galani, K., Mantero, J., et al. (2024). Single-cell multiregion dissection of Alzheimer’s disease. Nature 632, 858–868.

4. Mathys, H., Peng, Z., Boix, C.A., Victor, M.B., Leary, N., Babu, S., Abdelhady, G., Jiang, X., Ng, A.P., Ghafari, K., et al. (2023). Single-cell atlas reveals correlates of high cognitive function, dementia, and resilience to Alzheimer’s disease pathology. Cell 186, 4365–4385.e27.

5. Mathys, H., Davila-Velderrain, J., Peng, Z., Gao, F., Mohammadi, S., Young, J.Z., Menon, M., He, L., Abdurrob, F., Jiang, X., et al. (2019). Single-cell transcriptomic analysis of Alzheimer’s disease. Nature 570, 332–337.

6. Lau, S.-F., Cao, H., Fu, A.K.Y., and Ip, N.Y. (2020). Single-nucleus transcriptome analysis reveals dysregulation of angiogenic endothelial cells and neuroprotective glia in Alzheimer’s disease. Proc Natl Acad Sci U S A 117, 25800–25809.

7. Leng, K., Li, E., Eser, R., Piergies, A., Sit, R., Tan, M., Neff, N., Li, S.H., Rodriguez, R.D., Suemoto, C.K., et al. (2021). Molecular characterization of selectively vulnerable neurons in Alzheimer’s disease. Nat Neurosci 24, 276–287.

8. Gabitto, M.I., Travaglini, K.J., Rachleff, V.M., Kaplan, E.S., Long, B., Ariza, J., Ding, Y., Mahoney, J.T., Dee, N., Goldy, J., et al. (2024). Integrated multimodal cell atlas of Alzheimer’s disease. Nat. Neurosci. 27, 2366–2383.

9. Deng, Y., Bi, M., Delerue, F., Forrest, S.L., Chan, G., van der Hoven, J., van Hummel, A., Feiten, A.F., Lee, S., Martinez-Valbuena, I., et al. (2022). Loss of LAMP5 interneurons drives neuronal network dysfunction in Alzheimer’s disease. Acta Neuropathol. 144, 637–650.

10. Cain, A., Taga, M., McCabe, C., Green, G.S., Hekselman, I., White, C.C., Lee, D.I., Gaur, P., Rozenblatt-Rosen, O., Zhang, F., et al. (2023). Multicellular communities are perturbed in the aging human brain and Alzheimer’s disease. Nat Neurosci 26, 1267–1280.

11. Green, G.S., Fujita, M., Yang, H.-S., Taga, M., Cain, A., McCabe, C., Comandante-Lou, N., White, C.C., Schmidtner, A.K., Zeng, L., et al. (2024). Cellular communities reveal trajectories of brain ageing and Alzheimer’s disease. Nature 633, 634–645.

12. Miyoshi, E., Morabito, S., Henningfield, C.M., Das, S., Rahimzadeh, N., Shabestari, S.K., Michael, N., Emerson, N., Reese, F., Shi, Z., et al. (2024). Spatial and single-nucleus transcriptomic analysis of genetic and sporadic forms of Alzheimer’s disease. Nat Genet 56, 2704–2717.

13. Lee, C.Y., Riffle, D., Xiong, Y., Momtaz, N., Lei, Y., Pariser, J.M., Sikdar, D., Hwang, A., Duan, Z., and Zhang, J. (2024). Characterizing dysregulations via cell-cell communications in Alzheimer’s brains using single-cell transcriptomes. BMC Neurosci 25, 24.

14. Liu, A., Fernandes, B.S., Citu, C., and Zhao, Z. (2024). Unraveling the intercellular communication disruption and key pathways in Alzheimer’s disease: an integrative study of single-nucleus transcriptomes and genetic association. Alzheimers Res Ther 16, 3.

15. Vieth, B., Parekh, S., Ziegenhain, C., Enard, W., and Hellmann, I. (2019). A systematic evaluation of single cell RNA-seq analysis pipelines. Nature Communications 10, 4667.

16. Germain, P.-L., Sonrel, A., and Robinson, M.D. (2020). pipeComp, a general framework for the evaluation of computational pipelines, reveals performant single cell RNA-seq preprocessing tools. Genome Biol 21, 227.

17. Smith, R., Strandberg, O., Mattsson-Carlgren, N., Leuzy, A., Palmqvist, S., Pontecorvo, M.J., Devous, M.D., Ossenkoppele, R., and Hansson, O. (2020). The accumulation rate of tau aggregates is higher in females and younger amyloid-positive subjects. Brain 143, 3805–3815.

18. Boccalini, C., Peretti, D.E., Scheffler, M., Mu, L., Griffa, A., Testart, N., Allali, G., Prior, J.O., Ashton, N.J., Zetterberg, H., et al. (2025). Sex differences in the association of Alzheimer’s disease biomarkers and cognition in a multicenter memory clinic study. Alzheimers Res Ther 17, 46.

19. Wu, D., Bi, X., and Chow, K.H.-M. (2024). Identification of female-enriched and disease-associated microglia (FDAMic) contributes to sexual dimorphism in late-onset Alzheimer’s disease. J Neuroinflammation 21, 1.

20. Vila-Castelar, C., Akinci, M., Palpatzis, E., Aguilar-Dominguez, P., Operto, G., Kollmorgen, G., Quijano-Rubio, C., Blennow, K., Zetterberg, H., Falcon, C., et al. (2025). Sex/gender effects of glial reactivity on preclinical Alzheimer’s disease pathology. Mol Psychiatry 30, 1430–1439.

21. Zhou, Y., Song, W.M., Andhey, P.S., Swain, A., Levy, T., Miller, K.R., Poliani, P.L., Cominelli, M., Grover, S., Gilfillan, S., et al. (2020). Human and mouse single-nucleus transcriptomics reveal TREM2-dependent and TREM2-independent cellular responses in Alzheimer’s disease. Nat Med 26, 131–142.

22. Otero-Garcia, M., Mahajani, S.U., Wakhloo, D., Tang, W., Xue, Y.-Q., Morabito, S., Pan, J., Oberhauser, J., Madira, A.E., Shakouri, T., et al. (2022). Molecular signatures underlying neurofibrillary tangle susceptibility in Alzheimer’s disease. Neuron 110, 2929–2948.e8.

23. Palop, J.J., and Mucke, L. (2016). Network abnormalities and interneuron dysfunction in Alzheimer disease. Nat. Rev. Neurosci. 17, 777–792.

24. Bartas, K., Nguyen, M., Zhao, W., Hui, M., Nie, Q., and Beier, K.T. (2025). Analysis of changes in intercellular communications in Alzheimer’s disease reveals conserved changes in glutamatergic transmission in mice and humans. Sci Rep 15, 26248.

25. Dharshini, S.A.P., Sanz-Ros, J., Pan, J., Tang, W., Vallejo, K., Liu, Y.C., Otero-Garcia, M., and Cobos, I. (2026). Molecular signatures of resilience to Alzheimer’s disease in neocortical layer 4 neurons. Nat Commun 17. 10.1038/s41467-026-68920-4.

26. Saha, O., Melo de Farias, A.R., Pelletier, A., Siedlecki-Wullich, D., Landeira, B.S., Gadaut, J., Carrier, A., Vreulx, A.-C., Guyot, K., Shen, Y., et al. (2024). The Alzheimer’s disease risk gene BIN1 regulates activity-dependent gene expression in human-induced glutamatergic neurons. Mol Psychiatry 29, 2634–2646.

27. Lauterborn, J.C., Scaduto, P., Cox, C.D., Schulmann, A., Lynch, G., Gall, C.M., Keene, C.D., and Limon, A. (2021). Increased excitatory to inhibitory synaptic ratio in parietal cortex samples from individuals with Alzheimer’s disease. Nat Commun 12, 2603.

28. Chen, K.S., Noureldein, M.H., Rigan, D.M., Hayes, J.M., Loi, K.J., Hur, J., Savelieff, M.G., and Feldman, E.L. (2025). Interneuron transcriptomics reveals pathologic markers of Alzheimer’s disease progression. Neurobiol Dis 213, 107012.

29. Qu, W., Lam, M., McInvale, J.J., Mares, J.A., Kwon, S., Humala, N., Mahajan, A., Nguyen, T., Jakubiak, K.A., Mun, J.-Y., et al. (2024). Xenografted human iPSC-derived neurons with the familial Alzheimer’s disease APP mutation reveal dysregulated transcriptome signatures linked to synaptic function and implicate LINGO2 as a disease signaling mediator. Acta Neuropathol 147, 107.

30. Gautier, H.O.B., Evans, K.A., Volbracht, K., James, R., Sitnikov, S., Lundgaard, I., James, F., Lao-Peregrin, C., Reynolds, R., Franklin, R.J.M., et al. (2015). Neuronal activity regulates remyelination via glutamate signalling to oligodendrocyte progenitors. Nature Communications 6, 8518.

31. Moura, D.M.S., Brennan, E.J., Brock, R., and Cocas, L.A. (2022). Neuron to Oligodendrocyte Precursor Cell Synapses: Protagonists in Oligodendrocyte Development and Myelination, and Targets for Therapeutics. Front. Neurosci. 15, 779125.

32. Buchanan, J., da Costa, N.M., and Cheadle, L. (2023). Emerging roles of oligodendrocyte precursor cells in neural circuit development and remodeling. Trends Neurosci. 46, 628–639.

33. Pichet Binette, A., Gaiteri, C., Wennström, M., Kumar, A., Hristovska, I., Spotorno, N., Salvadó, G., Strandberg, O., Mathys, H., Tsai, L.-H., et al. (2024). Proteomic changes in Alzheimer’s disease associated with progressive Aβ plaque and tau tangle pathologies. Nature Neuroscience 27, 1880–1891.

34. Balcomb, K., Johnston, C., Kavanagh, T., Leitner, D., Schneider, J., Halliday, G., Wisniewski, T., Sunde, M., and Drummond, E. (2024). SMOC1 colocalizes with Alzheimer’s disease neuropathology and delays Aβ aggregation. Acta Neuropathol 148, 72.

35. Johnson, E.C.B., Bian, S., Haque, R.U., Carter, E.K., Watson, C.M., Gordon, B.A., Ping, L., Duong, D.M., Epstein, M.P., McDade, E., et al. (2023). Cerebrospinal fluid proteomics define the natural history of autosomal dominant Alzheimer’s disease. Nat Med 29, 1979–1988.

36. Zhang, P., Kishimoto, Y., Grammatikakis, I., Gottimukkala, K., Cutler, R.G., Zhang, S., Abdelmohsen, K., Bohr, V.A., Misra Sen, J., Gorospe, M., et al. (2019). Senolytic therapy alleviates Aβ-associated oligodendrocyte progenitor cell senescence and cognitive deficits in an Alzheimer’s disease model. Nature Neuroscience 22, 719–728.

37. Wang, C., Zong, S., Cui, X., Wang, X., Wu, S., Wang, L., Liu, Y., and Lu, Z. (2023). The effects of microglia-associated neuroinflammation on Alzheimer’s disease. Front. Immunol. 14, 1117172.

38. Mancuso, R., Fattorelli, N., Martinez-Muriana, A., Davis, E., Wolfs, L., Van Den Daele, J., Geric, I., Premereur, J., Polanco, P., Bijnens, B., et al. (2024). Xenografted human microglia display diverse transcriptomic states in response to Alzheimer’s disease-related amyloid-β pathology. Nat Neurosci 27, 886–900.

39. Nguyen, P.T., Dorman, L.C., Pan, S., Vainchtein, I.D., Han, R.T., Nakao-Inoue, H., Taloma, S.E., Barron, J.J., Molofsky, A.B., Kheirbek, M.A., et al. (2020). Microglial Remodeling of the Extracellular Matrix Promotes Synapse Plasticity. Cell 182, 388–403.e15.

40. Basilico, B., Ferrucci, L., Ratano, P., Golia, M.T., Grimaldi, A., Rosito, M., Ferretti, V., Reverte, I., Sanchini, C., Marrone, M.C., et al. (2022). Microglia control glutamatergic synapses in the adult mouse hippocampus. Glia 70, 173–195.

41. Parkhurst, C.N., Yang, G., Ninan, I., Savas, J.N., Yates, J.R., 3rd, Lafaille, J.J., Hempstead, B.L., Littman, D.R., and Gan, W.-B. (2013). Microglia promote learning-dependent synapse formation through brain-derived neurotrophic factor. Cell 155, 1596–1609.

42. Johnson, E.C.B., Carter, E.K., Dammer, E.B., Duong, D.M., Gerasimov, E.S., Liu, Y., Liu, J., Betarbet, R., Ping, L., Yin, L., et al. (2022). Large-scale deep multi-layer analysis of Alzheimer’s disease brain reveals strong proteomic disease-related changes not observed at the RNA level. Nature Neuroscience 25, 213–225.

43. Mostafavi, S., Gaiteri, C., Sullivan, S.E., White, C.C., Tasaki, S., Xu, J., Taga, M., Klein, H.-U., Patrick, E., Komashko, V., et al. (2018). A molecular network of the aging human brain provides insights into the pathology and cognitive decline of Alzheimer’s disease. Nature Neuroscience 21, 811–819.

44. Krzak, M., Raykov, Y., Boukouvalas, A., Cutillo, L., and Angelini, C. (2019). Benchmark and Parameter Sensitivity Analysis of Single-Cell RNA Sequencing Clustering Methods. Front Genet 10, 1253.

45. Hu, Q., and Greene, C.S. (2019). Parameter tuning is a key part of dimensionality reduction via deep variational autoencoders for single cell RNA transcriptomics. Pac Symp Biocomput 24, 362–373.

46. Luecken, M.D., Gigante, S., Burkhardt, D.B., Cannoodt, R., Strobl, D.C., Markov, N.S., Zappia, L., Palla, G., Lewis, W., Dimitrov, D., et al. (2025). Defining and benchmarking open problems in single-cell analysis. Nat Biotechnol 43, 1035–1040.

47. Raimundo, F., Vallot, C., and Vert, J.-P. (2020). Tuning parameters of dimensionality reduction methods for single-cell RNA-seq analysis. Genome Biol 21, 212.

48. Pullin, J.M., and McCarthy, D.J. (2024). A comparison of marker gene selection methods for single-cell RNA sequencing data. Genome Biology 25, 56.

49. Squair, J.W., Gautier, M., Kathe, C., Anderson, M.A., James, N.D., Hutson, T.H., Hudelle, R., Qaiser, T., Matson, K.J.E., Barraud, Q., et al. (2021). Confronting false discoveries in single-cell differential expression. Nat Commun 12, 5692.

50. Mukamel, E.A., and Yu, Z. (2025). False positives in study of memory-related gene expression. Nature 642, E1–E3.

51. Baran, Y., Bercovich, A., Sebe-Pedros, A., Lubling, Y., Giladi, A., Chomsky, E., Meir, Z., Hoichman, M., Lifshitz, A., and Tanay, A. (2019). MetaCell: analysis of single-cell RNA-seq data using K-nn graph partitions. Genome Biol 20, 206.

52. Bilous, M., Hérault, L., Gabriel, A.A., Teleman, M., and Gfeller, D. (2024). Building and analyzing metacells in single-cell genomics data. Mol Syst Biol 20, 744–766.

53. Liu, P., and Li, J.J. (2025). mcRigor: a statistical method to enhance the rigor of metacell partitioning in single-cell data analysis. Nat Commun 16, 8602.

54. Zhang, H., Tahami Monfared, A.A., Zhang, Q., and Honig, L.S. (2025). Incidence and Prevalence of Alzheimer’s Disease in Medicare Beneficiaries. Neurol Ther 14, 319–333.

55. Ferretti, M.T., Iulita, M.F., Cavedo, E., Chiesa, P.A., Schumacher Dimech, A., Santuccione Chadha, A., Baracchi, F., Girouard, H., Misoch, S., Giacobini, E., et al. (2018). Sex differences in Alzheimer disease - the gateway to precision medicine. Nat Rev Neurol 14, 457–469.

56. Emrani, S., and Sundermann, E.E. (2025). Sex/gender differences in the clinical trajectory of Alzheimer’s disease: Insights into diagnosis and cognitive reserve. Front Neuroendocrinol 77, 101184.

57. Coughlan, G.T., Klinger, H.M., Boyle, R., Betthauser, T.J., Binette, A.P., Christenson, L., Chadwick, T., Hansson, O., Harrison, T.M., Healy, B., et al. (2025). Sex Differences in Longitudinal Tau-PET in Preclinical Alzheimer Disease: A Meta-Analysis. JAMA Neurol 82, 364–375.

58. Belonwu, S.A., Li, Y., Bunis, D., Rao, A.A., Solsberg, C.W., Tang, A., Fragiadakis, G.K., Dubal, D.B., Oskotsky, T., and Sirota, M. (2022). Sex-Stratified Single-Cell RNA-Seq Analysis Identifies Sex-Specific and Cell Type-Specific Transcriptional Responses in Alzheimer’s Disease Across Two Brain Regions. Mol Neurobiol 59, 276–293.

59. Soudy, M., Bars, S.L., and Glaab, E. (2025). Sex-dependent molecular landscape of Alzheimer’s disease revealed by large-scale single-cell transcriptomics. Alzheimers Dement 21, e14476.

60. Lopez-Lee, C., Kodama, L., Fan, L., Zhu, D., Zhu, J., Wong, M.Y., Ye, P., Norman, K., Foxe, N.R., Ijaz, L., et al. (2024). Tlr7 drives sex differences in age-and Alzheimer’s disease–related demyelination. Science. 10.1126/science.adk7844.

61. Bliese, S.R., Basu, B., Beyer, S.E., Elsadany, M., Michaelson, J.J., and Chatterjee, S. (2025). Single-cell resolution spatial transcriptomic signature of the retrosplenial cortex during memory consolidation. Molecular Psychiatry 31, 1845–1856.

62. Olney, K.C., Gibson, K.A., Cadiz, M.P., Rahimzadeh, N., Swarup, V., and Fryer, J.D. (2025). Postmortem interval leads to loss of disease-specific signatures in brain tissue. eNeuro 12, ENEURO.0505–0524.2025.

63. Bennett, D.A., Wilson, R.S., Schneider, J.A., Evans, D.A., Aggarwal, N.T., Arnold, S.E., Cochran, E.J., Berry-Kravis, E., and Bienias, J.L. (2003). Apolipoprotein E epsilon4 allele, AD pathology, and the clinical expression of Alzheimer’s disease. Neurology 60, 246–252.

64. Wolf, F.A., Angerer, P., and Theis, F.J. (2018). SCANPY: large-scale single-cell gene expression data analysis. Genome Biol 19, 15.

65. Kim, J.K., Kolodziejczyk, A.A., Ilicic, T., Teichmann, S.A., and Marioni, J.C. (2015). Characterizing noise structure in single-cell RNA-seq distinguishes genuine from technical stochastic allelic expression. Nat Commun 6, 8687.

66. Kharchenko, P.V., Silberstein, L., and Scadden, D.T. (2014). Bayesian approach to single-cell differential expression analysis. Nat Methods 11, 740–742.

67. Servén, D., Brummitt, C., Király, F., JodesL, Schmitz, D., Carmichael, Z., Ouslan, A., Miccoli, F., IlyesD, Taylor, J., et al. (2025). dswah/pyGAM: v0.12.0 (Zenodo) 10.5281/ZENODO.1208723.

68. Troulé, K., Petryszak, R., Cakir, B., Cranley, J., Harasty, A., Prete, M., Tuong, Z.K., Teichmann, S.A., Garcia-Alonso, L., and Vento-Tormo, R. (2025). CellPhoneDB v5: inferring cell-cell communication from single-cell multiomics data. Nat Protoc. 10.1038/s41596-024-01137-1.

69. Fang, Z.F., Liu, X., and Peltz, G. (2023). GSEApy: a comprehensive package for performing gene set enrichment analysis in Python. Bioinformatics 39. 10.1093/bioinformatics/btac757.

70. Oveisgharan, S., Yang, J., Yu, L., Burba, D., Bang, W., Tasaki, S., Grodstein, F., Wang, Y., Zhao, J., De Jager, P.L., et al. (2023). Estrogen Receptor Genes, Cognitive Decline, and Alzheimer Disease. Neurology 100, e1474–e1487.

71. Vallat, R. (2018). Pingouin: statistics in Python. Journal of Open Source Software 3, 1026.

